# A chemical strategy to control protein networks in vivo

**DOI:** 10.1101/2020.04.08.031427

**Authors:** Michael J. Ziegler, Klaus Yserentant, Volker Middel, Valentin Dunsing, Antoni J. Gralak, Kaisa Pakari, Jörn Bargstedt, Christoph Kern, Salvatore Chiantia, Uwe Strähle, Dirk-Peter Herten, Richard Wombacher

## Abstract

Direct control of protein interaction by chemically induced protein proximity (CIPP) holds great potential for cell- and synthetic biology as well as therapeutic applications. However, toxicity, low cell-permeability and lack of orthogonality currently limit the use of available chemical inducers of proximity (CIP). We present ‘Mandi’, a novel CIP and demonstrate its applicability in cell culture systems as well as living organisms for protein translocation, protein network shuttling and manipulation of endogenous proteins.

## MAIN TEXT

Protein proximity is a key regulatory mechanism in cellular processes, including metabolic pathways and cellular signaling, which are essential to sustain cellular integrity and to organize cellular response. Tools to investigate and manipulate protein proximity must meet a range of demanding requirements such as fast dose response, high efficiency and spatial control. At the same time, they should neither interfere with the process under study or off-target cellular processes nor be cytotoxic. Chemical inducers of proximity (CIP) are small, drug-like molecules that induce protein proximity by mediating interactions between specific receptor and receiver domains and have been widely used in biology (Fig. 1a)^1^. Different chemically induced protein proximity (CIPP) systems have been successfully applied for induced signal transduction^2–4^, transcription control^5,6^, protein translocation^7^, degradation^8^, aggregation^9^ or regulation of chromatin^10,11^. Therefore they hold great potential for future drug development by specific control of metabolic pathways and signaling cascades^1^. Over the past few years, phytohormone based CIPP systems have received significant attention since they make use of plant proteins, which do not occur in the animal kingdom and are therefore fully orthogonal to processes in mammalian cells. Gibberellic acid (GA_3_) as well as abscisic acid (ABA) induce protein-protein interactions upon ligand binding to regulate plant growth^12^ or stress resistance^13^. Both, GA_3_ and ABA in combination with their dimerization domains Gibberellin-insensitive dwarf protein 1/Gibberellic acid-insensitive (GID1/GAI) and Pyrabactin resistance like (PYL)/Abscisic Acid insensitive (ABI) respectively, have been used as CIPP systems with times-to-effect in the range of minutes^14,15^. Recently, engineered ABA receptors have been reported for agrochemical control of water use in plants^16^. The genetically modified receptors do not respond to the phytohormone ABA but to the agrochemical mandipropamid (Mandi), a fungicide extensively used in agriculture (Fig. 1b). A sextuple mutant PYR^Mandi^ of the ABA receptor Pyrabactin resistance 1 (PYR1) was identified that specifically binds mandipropamid^16^ replacing the natural ABA response in plants (Supplementary Fig. 1). We hypothesized that, like the GA_3_ and ABA systems, Mandi and the respective receptor PYR^Mandi^ can be used as a CIP in mammalian cells. With its simple molecular structure, Mandi is readily available either by chemical synthesis^17^ or commercially as pure compound. We therefore propose Mandi as an attractive candidate to overcome current limitations of CIPP systems to leverage these tools for in vivo applications.

**Fig. 1:**
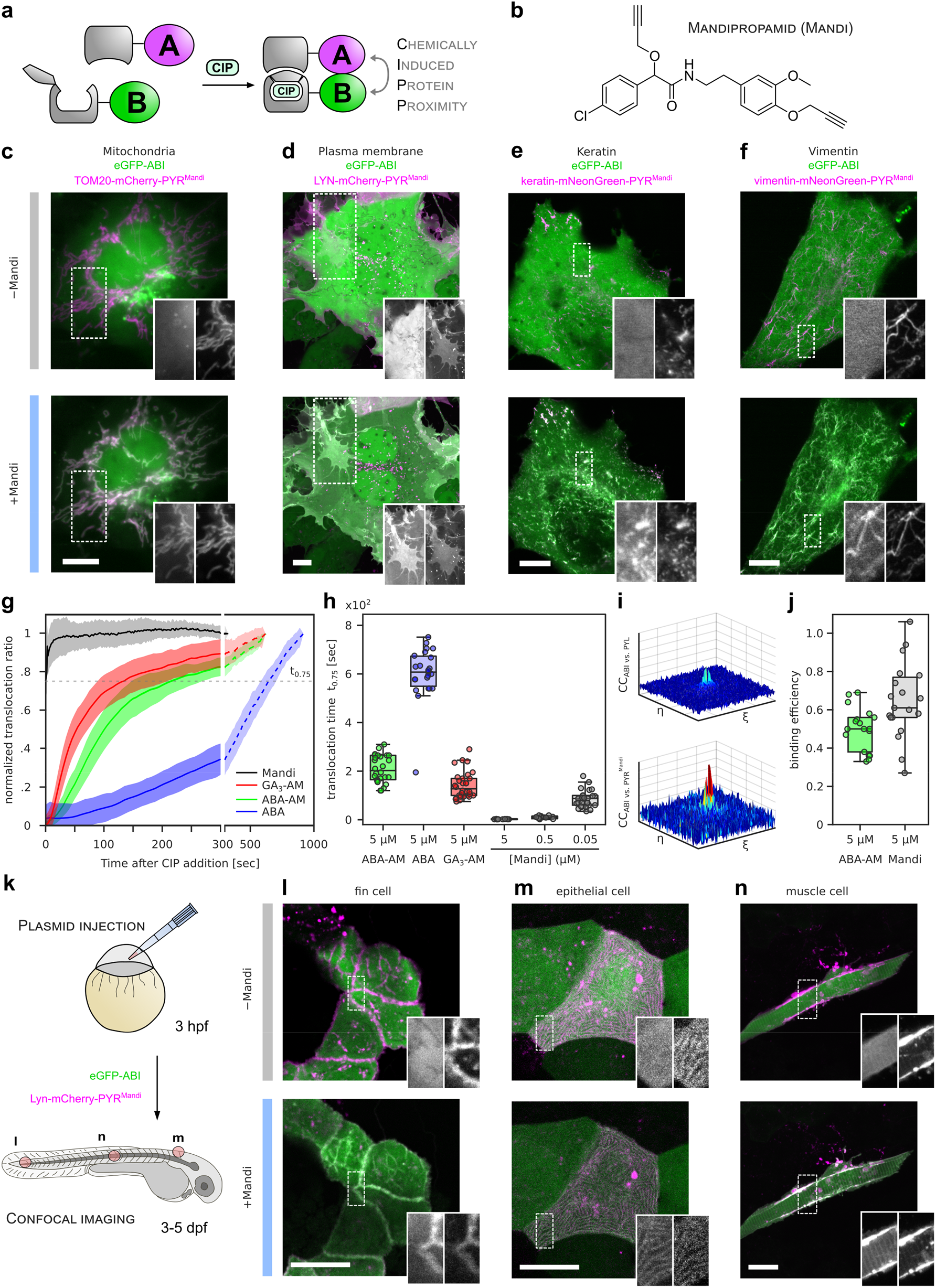
Mandi - a new and ultrafast chemical inducer of proximity (CIP). **a,** Chemically induced protein proximity (CIPP) to control protein-protein interaction between two proteins of interest A and B. **b,** Chemical structure of mandipropamid (Mandi). **c,** Live-cell epifluorescence microscopy images of COS-7 cells transiently transfected with TOM20-mCherry-PYR^Mandi^-IRES-eGFP-ABI before and 5 min after addition of Mandi (100 nM). Colocalization completed within 1 min (Supplementary video 1). **d**, Confocal live-cell microscopy images of COS-7 cells co-transfected with LYN-mCherry-PYR^Mandi^ and eGFP-ABI before and 2 min after addition of 100 nM Mandi. **e,f,** Confocal live-cell microscopy images of COS-7 cells transfected with Keratin **(e)**/Vimentin**(f)**-mNeonGreen-PYR^Mandi^-IRES-Halo-ABI. **c-f**, Halo-ABI was labeled with HTL-SiR. Images were acquired before and 5 min after addition of 50 nM Mandi. Representative data for 6-20 cells in 1-2 independent experiments, Scale bar: 10 μm. **g,** Ratio of cytosolic receiver domain localized in cytosol and on mitochondria over time. Data are normalized to ratio before addition of CIP and after translocation was completed. Mean (line) ±1 SD (shaded region) for each condition from 11 (Mandi, n=2 independent experiments), 30 (GA_3_-AM, n=4), 30 (ABA-AM, n=4) and 23 (ABA, n=4) cells. Translocation time t_0.75_, *i.e.* time at which translocation to mitochondria was 75 % of maximum, shown as dashed gray line. **h,** t_0.75_, for ABA, ABA-AM and GA_3_-AM (5 μM, respectively) and Mandi (5, 0.5, 0.05 μM) CIP systems shown in g. Each dot represents t_0.75_ from a single cell. Box plots indicate median, 25^th^ and 75^th^ percentile for each condition. Whiskers extend to 1x interquartile range. Mandi datasets at varying concentration with 11 (n=2 independent experiments), 13 (n=2), 29 (n=4) cells per condition. **i**, Representative cross-correlation functions for mCherry-ABI vs. eGFP-PYL (top) and mCherry-ABI vs. YFP-PYR^Mandi^ (bottom) obtained from RSICS measurements in transiently transfected COS-7 cells after addition of 5 μM ABA-AM or 5 μM Mandi. **j**, Binding efficiencies obtained from RSICS relative cross-correlation amplitudes (see methods) for mCherry-ABI binding to eGFP-PYL after ABA-AM addition or to PYR^Mandi^ after Mandi addition. Data from 18 (ABA-AM) or 19 (Mandi) cells pooled from 2 independent experiments per condition. Box plots as in h. **k**, Schematic illustration of workflow for in vivo application in zebrafish embryos. Fertilized eggs were injected with vectors for Lyn-or TOM20-mCherry-PYR^Mandi^ (Supplementary Fig. 10) and eGFP-ABI expression resulting in mosaic expression of target proteins at 3-5 dpf. **l, m, n,** Confocal microscopy images of different cell types in living zebrafish embryos expressing receiver and plasma membrane localized receptor domains before and 10-20 min after addition of 500 nM Mandi. Representative data from ≥3 independent experiments for each cell type. Scale bar: 40 μm.

To test if Mandi can induce protein proximity in mammalian cells, we established a reporter assay based on colocalization of fluorescently labeled fusion proteins (Supplementary Fig. 2). We expressed the receptor domain PYR^Mandi^ fused to different intracellular proteins with characteristic localization and the receiver domain ABI as a cytosolic protein. Addition of Mandi resulted in rapid colocalization at the respective targets in all tested cell lines (Fig. 1c-f, 2b, Supplementary Fig. 3,10,15). While addition of Mandi to a final concentration of 1 μM resulted in efficient colocalization within seconds, colocalization using 100 nM Mandi was completed within 1 min. At 10 nM, colocalization was still detectable after 4 min, though less efficient (Supplementary Fig. 3, Supplementary video 1,2). To quantitatively show the superior performance of Mandi based CIPP over existing approaches, we performed a direct comparison with other phytohormone based CIPP systems (Supplementary Fig. 4-7). Since GA_3_ and ABA exhibit no or low membrane permeability, we used acetoxymethyl (AM) ester-modified derivatives with improved membrane permeability for both^16,18^. To determine the time-to-effect for each CIP, we extracted the recruitment kinetics for a cytosolic receiver to its corresponding receptor domain targeted to the outer mitochondrial membrane using a TOM20 fusion protein. Using an automated epifluorescence microscopy platform with integrated liquid handling, we performed time-lapse imaging after addition of CIP and used a machine-learning approach for automated cell segmentation and subsequent intensity readout^19,20^. While ABA, ABA-AM and GA_3_-AM at 5 μM show receiver recruitment to mitochondria as defined as the translocation ratio (t_0.75_, see methods) within 10±1.8, 3.5±0.9 and 2.4±0.9 min, respectively, the translocation induced by Mandi at the same concentration is too fast to be resolved by our assay (≤2.4 sec, Fig. 1g,h). At 100x lower Mandi concentration, translocation occurred within 1.4±0.6 min, which is still >1.7 times faster than with GA_3_-AM at 5 μM (Fig. 1h). In addition to determining the times-to-effect, we measured the relative amount of receiver bound to receptor in absence and presence of CIPs using raster spectral image correlation spectroscopy (RSICS)^21^. After transient expression of cytosolic receiver and receptor domains fused to spectrally different fluorescent proteins, we determined the interacting fraction by computing the cross-correlation functions (CCF) between spectral channels^22^ (see methods). We found that after addition of Mandi at 5 μM concentration, 65±20% of ABI and PYR^Mandi^ were bound in complexes, while after stimulation with ABA-AM at 5 μM concentration, only 50±11% of ABI and PYL were bound (Fig i,j). The 70-to ~300-fold faster induction of protein interaction compared to GA_3_-AM and ABA, respectively as well as the higher binding efficiency of Mandi compared to ABA-AM, shows the superior performance of Mandi and demonstrates its enormous potential for live-cell and in vivo applications, an up to now unmet challenge for CIPP. To the best of our knowledge, GA_3_ and ABA have not been used in living organisms so far, presumably because both possess low membrane permeability and tissue penetration. The most widely used CIP rapamycin is both toxic and immunosuppressive and, consequently, of limited use for applications in living organisms due to its narrow therapeutic window^23^. As Mandi follows the rule of five and possesses drug-likeness while being non-toxic^24^, we hypothesized that it may be ideally suited for in vivo applications. To test this, we expressed receiver and receptor domains on various cellular targets in zebrafish (*Danio rerio*) embryos and evaluated Mandi’s ability to induce protein proximity in different tissues of 3-5 days post fertilization (dpf) embryos (Fig. 1k). At concentrations as low as 500 nM, Mandi successfully induced protein colocalization within minutes at subcellular targets, *i.e.* plasma membrane or mitochondria, in different tissues (Fig. 1l,m,n, Supplementary Fig. 10, Supplementary video 3). Remarkably, addition of Mandi solution on top of the agarose embedded embryos was sufficient to achieve colocalization in cells deep in the tissue (*e.g.* muscle cells) within minutes, reflecting its excellent tissue penetration. As expected based on risk assessments related to its use in agriculture^24^, we did not observe toxicity of Mandi in zebrafish embryos even at concentrations up to a 100 times above required working concentrations (Supplementary Fig. 11).

**Fig. 2:**
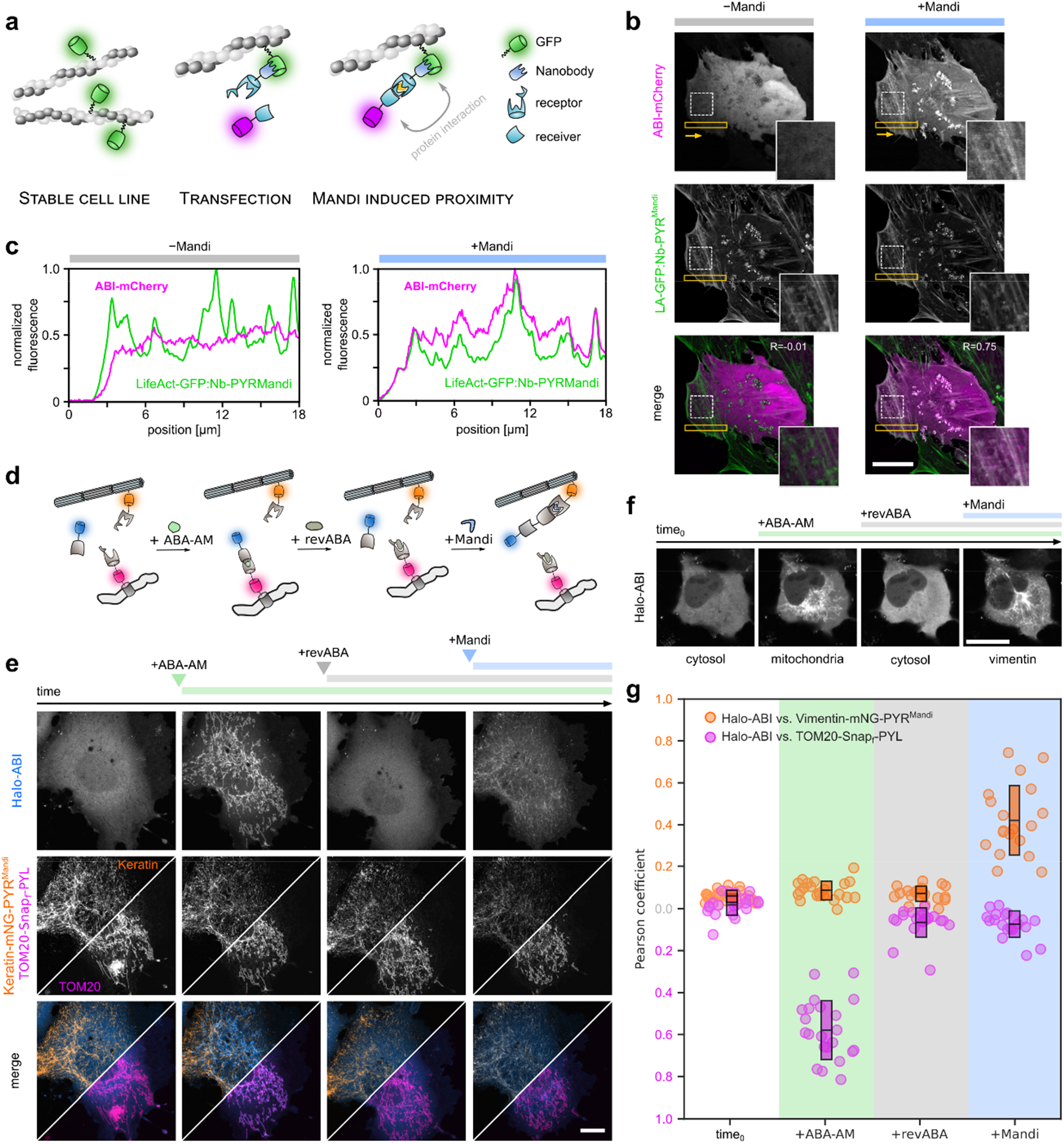
Nanobody assisted targeting of chemically induced protein proximity (natCIPP) and reversible and dynamic protein shuttling in living cells. **a**, Schematic illustration **(a)** and live-cell imaging **(b)** of natCIPP. HeLa cells stably expressing LifeAct-GFP were transfected with antiGFP-nanobody-PYR^Mandi^ and mCherry-ABI fusions. Images acquired before (Pearson’s R = 0.01) and 5 min after addition of 50 nM Mandi (Pearson’s R = 0.75). Representative data for 30 cells in 2 independent experiments. Scale bar: 20 μm. **c**, Line profiles of eGFP and mCherry signal intensity in ROI (yellow box) before and after addition of Mandi. **d**, Schematic illustration of four step procedure to shuttle cytosolic protein between different cellular targets. **e**, Confocal fluorescence microscopy images of shuttling process between keratin and mitochondria in a living cell. COS-7 cells were co-transfected with Keratin-mNeonGreen-PYR^Mandi^-IRES-Halo-ABI and TOM20-SNAP_f_-PYL. Halo-ABI and SNAP_f_-PYL were labelled with HTL-SiR and TMR-Star, respectively. Upper row shows dynamic receiver localization. Middle row shows receptor localizations as references. Split images depict keratin and mitochondrial localization in two different channels. Lower row are respective merges. Images acquired at t_0_, 10 min after addition of ABA-AM (200 nM), 25 min after addition of revABA (20 μM), 10 min after addition of Mandi (200 nM). Scale bar: 20 μm. **f,** Representative confocal fluorescence microscopy images of shuttling between vimentin and mitochondria over time, only dynamic receiver localization depicted (reference and merge illustrated in Supplementary Fig. 16). Scale bar: 20 μm. Representative data from 22 cells in 2 independent experiments. **g,** Pearson correlation coefficients between receiver and receptors at different time points for data set shown in f, indicating signal correlation quality. Boxes indicate mean ±1 SD for each condition.

Manipulation of endogenous proteins to enable protein interaction studies at native concentrations in their physiological environment is highly desirable. However, tagging of endogenous proteins can result in altered expression patterns and ill-defined perturbations of protein function. While small drug-like probes for specific protein manipulation have been shown to be highly useful in cell biology research^25^, a technology that can be applied to arbitrary native proteins is currently not available. Recent advances using nanobodies have shown great potential for endogenous protein targeting in living cells^26,27^. We hypothesized that nanobody assisted targeting in combination with the Mandi CIPP system could induce artificial interactions between endogenous proteins and any genetically introduced effector protein in a dynamic and controlled manner. As a proof of principle in living cells, we used a well-studied antiGFP-nanobody^28^ in combination with cell lines stably expressing F-actin binding LifeAct-GFP or paxillin-YFP. The antiGFP-nanobody and mCherry are expressed as fusion proteins with the Mandi receptor and receiver, respectively (Fig. 2a). The nanobody thus serves as an adaptor between the native target and the artificial CIPP system, placing the interaction of effector and endogenous target protein under strict control of Mandi. This is visualized by the appearance of characteristic structures upon addition of Mandi (Fig. 2b,c, Supplementary Fig. 12). Such nanobody assisted targeting of chemically induced protein proximity (natCIPP) can easily be extended to other targets^29^.

The simultaneous use of multiple CIPP systems allows the construction of Boolean logic gates and enables the design of artificial genetic circuits^15,30^. For such applications, the CIPP systems must be orthogonal to the organism under study and among themselves. We tested if the Mandi system could be used in conjunction with GA_3_ and ABA CIPP systems to create complex logic gates in cell culture systems. As expected, we found Mandi to be fully orthogonal to GA_3_ (Supplementary Fig. 13). ABA and Mandi, however, recruit the identical receiver domain ABI and are therefore semi-orthogonal (Supplementary Fig. 14). We further confirmed the compatibility of the ABA and Mandi CIPP systems by simultaneously expressing both systems and measuring the interacting fractions before CIP addition and after ABA-AM as well as Mandi addition using RSICS (Supplementary Fig. 9d). Semi-orthogonal CIPP systems with a single receiver and multiple receptors represent a powerful tool for advanced applications, previously only achievable using double fusion constructs^31^.

A major challenge in synthetic biology is to mimic complex and highly dynamic intracellular protein networks and to further manipulate their regulation through external stimuli. We designed a protein translocation system based on semi-orthogonal CIPPs where a cytosolic receiver protein is reversibly shuttled between different intracellular targets depending on the specific CIP input (Fig. 2d). Such applications are limited by competing interactions between multiple receptors and the receiver^32^. We addressed this problem by making use of an ABA antagonist, revABA (Supplementary Fig. 15a), previously used in crop science^33^. We used the antagonist as a suppressive stimulus to inhibit one of the interactions within the network (Supplementary Fig. 15b,c, Supplementary video 4). Subsequent addition of ABA-AM, revABA and Mandi was then used to shuttle the cytosolic receiver between different subcellular localizations (Fig 2e). Intracellular shuttling could be performed between different targets, in different cell types (Fig. 2f, Supplementary Fig. 16, 17), and was highly efficient (Fig. 2g).

In summary, we demonstrate that Mandi based CIPP is a versatile technology to reversibly control the localization and interaction of proteins within cellular networks, not only in living cells, but also in the more complex context of living organisms. In combination with nanobodies, Mandi allows targeting of endogenous, untagged target proteins using natCIPP. The excellent cell permeability and low toxicity of Mandi enable applications at minimal concentrations in cultured cells and in vivo. Based on these findings, we expect Mandi CIPP technology to become a powerful tool for manipulating protein localization and interaction in cell biological research, as well as for circuit design in synthetic biology. Furthermore, we believe Mandi to hold great potential for future use in therapeutic applications such as PROTAC approaches or in CAR-T cell therapies^34,35^.

## ACKNOWLEDGEMENTS

R.W. acknowledges funding from the Deutsche Forschungsgemeinschaft DFG (SPP1623, WO 1888/1-2) and D.-P.H. from the Federal Ministry of Education and Research (BMBF/VDI; MorphiQuant3D and Switch-Click-Microscopy). M.J.Z gratefully acknowledges fellowship by the Carl-Zeiss-Stiftung. We thank Steven Thomas (College of Medical and Dental Sciences, University of Birmingham, UK) for helpful discussion about our work and the manuscript. We thank Ada E. Cavalcanti-Adam for REF cells stably expressing paxilin-YFP and Jacob Piehler for HeLa cells stably expressing LifeAct-GFP-Halo. We thank Josef Denzer for providing Revus Top^®^. We gratefully acknowledge access to the Nikon Imaging Center at Heidelberg University.

## AUTHOR CONTRIBUTIONS

M.J.Z, K.Y. and R.W. designed this study. A.G. and M.J.Z. designed and cloned vectors. C.K. and J.B. synthesized revABA. M.J.Z. synthesized Mandi and ABA-AM. K.Y. designed and established software for automated microscopy and data analysis. K.Y., K.P and M.J.Z. performed live cell microscopy. K.Y. analyzed microscopy data. V.D. and S.C. planned, performed and analyzed RSICS experiments. V.M. performed in vivo studies. M.J.Z., K.Y. and R.W. analyzed data. D.P.H, U.S., S.C. and R.W. supervised research. M.J.Z., K.Y and R.W wrote the manuscript with support from all authors.

## COMPETING INTERESTS

The authors declare no competing interests.

## Notes

https://www.dropbox.com/s/9koqn06jc6pfpqq/Ziegleretal_SM1.avi?dl=0

https://www.dropbox.com/s/u5ktmd9brbyjxve/Ziegleretal_SM2.avi?dl=0

https://www.dropbox.com/s/464vuloc0w40nhr/Ziegleretal_SM3.avi?dl=0

https://www.dropbox.com/s/k59lcxy12rixehj/Ziegleretal_SM4.avi?dl=0

