## Supporting Information for "A chemical strategy to control protein networks in vivo"


1 CONSTRUCT DESIGN AND PREPARATION 4

**1.1 General procedure 4**

**1.2 Primer List 4**

**1.3 Plasmids 6**

2 CELL CULTURE, TRANSFECTION AND SAMPLE PREPARATION 9

**2.1 General remarks 9**

**2.2 Protein shuttling and co-localization analysis 9**

**2.3 Nanobody assisted targeting of chemically induced protein proximity (natCIP) 10**

3 WIDEFIELD EPIFLUORESCENCE MICROSCOPY 10

4 CONFOCAL FLUORESCENCE MICROSCOPY 10

5 AUTOMATED SCREENING OF CIP EFFICIENCY 11

6 RASTER SPECTRAL IMAGE CORRELATION SPECTROSCOPY (RSICS) 13

7 IN VIVO EXPERIMENTS 15

**7.1 Zebrafish strains 15**

**7.2 Real-time imaging of zebrafish embryos 15**

8 SUPPLEMENTARY FIGURES 16

9 SUPPLEMENTARY VIDEOS 31

10 MATERIALS 33

**10.1 Synthesis 33**

11 NMR CHARACTERIZATION 38

**11.1 Mandipropamid 38**

**11.2 ABA-AM 39**

**11.3 (p-Tolyl)prop-2-yn-1-ol 40**

**11.4 3-(p-Tolyl)prop-2-yn-1-yl 4-methylbenzolsulfonate 41**

**11.5 ABA-(*S*)-OH 42**

**11.6 revABA 43**

12 REFERENCES 44

### 1 Construct design and preparation

#### 1.1 General procedure

For construction of new plasmids, fragments were amplified by PCR from appropriate sources. Primers used for PCR (see Supplementary Table 1) delivered by Integrated DNA Technologies, Inc. (IDT, Belgium). PCR reaction mix of backbone fragments were digested with DPN1 (addition of 10 µl CutSmart Buffer + 1 µl DPN1 to 50 µl PCR mix, incubation at 37°C for 1h. All PCR fragments were purified by preparative agarose gel electrophoresis and extracted with QIAquick Gel Extraction Kit (Qiagen, Germany). Ligation by Gibson assembly was performed in equimolar ratio of all fragments. Plasmid sequence was validated by Sanger sequencing (Seqlab, Germany) using either standard primers or premixed sequence primers. All plasmids will be made available via Addgene after final publication of manuscript.

#### 1.2 Primer List

**Supplementary Table 1: Full list of all used primers.**

| Primer | Sequence (5'-3') |
|---|---|
| #1 | TAA ATG CAG AAC GGC CCC GGT GCT C |
| #2 | CAT GGT GGC GGC CGC AAT TGC GCT AG |
| #3 | TAA ACC GGA ATT CCG |
| #4 | CAT GGT TGT GGC CAT ATT ATC ATC |
| #5 | GCA ATT GCG GCC GCC ACC ATG GTG AGC AAG GGC GAG GAG CTG |
| #6 | GAG GGG CGG AAT TCC GGT TTA ACA TTC CGC GTT TAC AAA CGC |
| #7 | GAT AAT ATG GCC ACA ACC ATG GGT CGG AAC AGC GCC ATC |
| #8 | GCA ATT GCG GCC GCC ACC ATG GTG AGC AAG GGC GAG GAG |
| #9 | ACG CGT GTG CCT TTG TAT GGT |
| #10 | CAT GAA TTC CGA AAA TGG ATA |
| #11 | CCA ACT CAA GAC GAA TTC ACC |
| #12 | CAT GAG CTC GGC CAT ATT ATC |
| #13 | TAT CCA TTT TCG GAA TTC ATG CCA CCA TGG ATG GGT CGG AAC |
| #14 | GGT GAA TTC GTC TTG AGT TGG TCC TCC TGC GCT CTT GTA CAG |
| #15 | GAT AAT ATG GCC GAG CTC ATG GTG AGC AAG GGC GAG GAG CTG |
| #16 | ACC ATA CAA AGG CAC ACG CGT TCC TCC TGC GCT CTT GTA CAG |
| #17 | TGG TCC TCC TGC GCT CTT GTA CAG |
| #18 | TGA GGA TCC GCC CCT CTC CCT |
| #19 | GTG CCT TTG TAT GGT TTT AC |
| #20 | GAA GTT GGG GTC ACT TCG TC |
| #21 | CGA AGT GAC CCC AAC TTC AGC GCA GGA GGA CCA ATG |

| | |
|---|---|
| #22 | TCC GGA TCC CAT GAG CTC GGC CAT ATT ATC ATC C |
| #23 | GAG CTC ATG GGA TCC GGA GCA AGT GGA AT |
| #24 | AGT AAA ACC ATA CAA AGG CAC TGG TCC TCC TGC GCT CTT GT |
| #25 | CGA AGT GAC CCC AAC TTC AGC GCA GGA GGA CCA ATG C |
| #26 | TCG GAT CCC ATG AGC TCG GCC ATA TTA T |
| #27 | AGC TCA TGG GAT CCG AAA TCG GTA CTG G |
| #28 | AGT AAA ACC ATA CAA AGG CAC ACC GGA AAT CTC CAG AGT AG |
| #29 | GGT GAA TTC ACC GGT ACC TG |
| #30 | TGA GCG GCC GCA TAG ATA AC |
| #31 | GGT ACC GGT GAA TTC ACC ACT CAA GAC GAA TTC ACC CA |
| #32 | CTA TGC GGC CGC TCA GTT CAT AGC TTC AGT GAT CG |
| #33 | AGC GCA GGA GGA CCA ATG |
| #34 | CCA TGG TGG CAT GAA TTC CG |
| #35 | GAA TTC ATG CCA CCA TGG ATG TCC ACC AGG TCC GTG TC |
| #36 | TGG TCC TCC TGC GCT CTT GTA CAG CTC GTC CAT GC |
| #37 | GAA TTC ATG CCA CCA TGG ATG AGC TTC ACC ACT CGC TC |
| #38 | TGG TCC TCC TGC GCT CTT GTA CAG CTC GTC CAT GC |
| #39 | TCA CAC TGG CGG CCG TCT AGA ATG GGT CGG AAC AGC GCC ATC |
| #40 | TAT AGA ATA GGG CCC CTC GAG TCA AGC GTA ATC TGG AAC ATC |
| #41 | AGC GCA GGA GGA CCA ACT CA |
| #42 | TCC ACT TGC TCC GGA TCC GA |
| #43 | TCC GGA GCA AGT GGA ATG GAC AAA GAC TGC GAA ATG AA |
| #44 | TGG TCC TCC TGC GCT ATT AAC CTC GAG TTT AAA CGC GG |
| #45 | AGC GCA GGA GGA CCA ATG CC |
| #46 | CCA TGG TGG CAT GAA TTC CG |
| #47 | GAA TTC ATG CCA CCA TGG ATG CAG GTT CAA CTG GTG GA |
| #48 | TGG TCC TCC TGC GCT TTT AGA GCT CAC CGT CAC CTG |
| #49 | GAT TAC AAG GAC GAT GAC GAT AAG TGA AGC GGC |
| #50 | ATC CTC ACG TGA CCT GGC TGC CAG AGC CAT CAC CT |
| #51 | ATT CAT GCC ACC ATG GAT GGT GAG CAA GGG CGA GGA |
| #52 | CCA TGG TGG CAT GAA TTC CGA AAA TGG ATA TA |
| #53 | CTT GTA CAG CTC GTC CAT GC |
| #54 | CTG ATC ATA ATC AGC CAT ACC ACA |
| #55 | TGG ACG AGC TGT ACA AGA GCG CAG GAG GAA CGC GT |
| #56 | TGG TAT GGC TGA TTA TGA TCA GTC ACT TAT CGT CAT CGT CCT TGT |
| #57 | CAT GGT GGC GAC CGG TAG |
| #58 | CCG GTC GCC ACC ATG GGT CGG AAC AGC GCC ATC |

| | |
|---|---|
| #59 | TGG TAT GGC TGA TTA TGA TCA GTC ACG TGA CCT GGC TGC CAG A |
| #60 | GGA TCC GGA GCA AGT GGA AT |
| #61 | CCG GTC GCC ACC ATG GGA TGT ATT AAA TCA AAA AGG AAA GAC GGA TCC GGA GCA AGT |
| #62 | ACT TGC TCC GGA TCC GTC TTT CCT TTT TGA TTT AAT ACA TCC CAT GGT GGC GAC CGG |
| #63 | TGG TCC TCC TGC GCT CTT GTA CAG |
| #64 | TGA GGA TCC GCC CCT CTC CCT |
| #65 | CTT TAC AGC TCG TCC ATG CCG AG |
| #66 | ATT ACG CTT GAA GCG GCC GCG ACT CTA GAT |
| #67 | TGG ACG AGC TGT ACA AGG GAT CCG GAG CAA GTG GAA |
| #68 | GCC GCT TCA AGC GTA ATC TGG AAC ATC |
| #69 | AGC GCA GGA GGA CCA ATG C |
| #70 | CAT GGT GGC GAC CGG TAG |
| #71 | TGG TCC TCC TGC GCT CTT GTA TAG CTC GTC CAT GCC G |
| #72 | CCG GTC GCC ACC ATG GTG AGC AAG GGC GAG GAG |

#### 1.3 Plasmids

**EGFP-GID1-IRES-TOM20-mCherry-GAI** was cloned from pcDNA3-PLE-IRES-GFP-GID1, which was a gift from Aline Daniel (Heidelberg University). For that, PLE and GFP-GID1 were removed by amplification of pcDNA3 backbone and IRES site with flanked primers (#1, #2, #3, #4). PLE was substituted by EGFP-GID1 which was amplified with appropriate overlaps from EGFP-GID1 (Addgene, #37306) (#5, #6). GFP-GID1 was substituted with TOM20-mCherry-GAI fragment amplified with appropriate overlaps from TOM20-mCherry-GAI plasmid (Addgene, #37306) (#7, #8). All fragments were ligated by Gibson Assembly and sequenced using standard primers (T7-fw, IRES-rev, SP6**).**

**TOM20-mCherry-PYL-IRES-EGFP-ABI** was generated from pSV-ABAactDA (#38247), TOM20-mCherry-GAI (#37316) and EGFP-GID1 (#37306) all obtained from Addgene. From pSV-ABAactDA, VP16AD and GAL4DBD were removed and exchange by TOM20-mCherry and EGFP. For this purpose, two backbone fragments were amplified by PCR using flanked primers (#9, #10, #11, #12). TOM20-mCherry-GAI delivered the TOM20-mCherry fragment (#13, #14) EGFP-GID1 originated the EGFP fragment (#15, #16). All fragments were ligated by Gibson Assembly and sequenced using standard primers (IRES-gap, SV-40 for, SV-40-pA-rev, IRES-rev).

**TOM20-mCherry-PYR$^{\mathbf{Mandi}}$-IRES-EGFP-ABI** was generated from TOM20-mCherry-PYL-IRES-EGFP-ABI. For that, the PYL side was removed by amplification of the backbone using flanked primers (#63, #64). PYR$^{\mathrm{Mandi}}$ gene fragment was delivered as gBlock (CTG TAC AAG AGC GCA GGA GGA CCA ATG CCA

TCT GAA TTG ACC CCT GAG GAA CGC TCC GAA TTG AAA AAT TCA ATC GCC GAA TTC CAT ACC TAT CAG CTC GAC CCC GGA TCT TGC AGT TCA CTG CAT GCA CAG CGC ATC CAC GCG CCC CCA GAA TTG GTG TGG TCT ATC GTT CGC CGC TTT GAC AAA CCC CAA ACG CAC CGG CAC TTC ATA AAG TCA TGT TCA GTT GAA CAG AAT TTC GAA ATG CGA GTG GGC TGC ACC AGA GAT ATA ATA GTA ATA TCC GGT CTC CCT GCA AAT ACA TCC ACG GAG CGA CTG GAC ATA CTT GAC GAT GAA AGA AGA GTT ACG GGC GCT TCT ATA ATT GGG GGC GAA CAC CGG CTG ACT AAC TAT AAG GGC GTC ACA ACC GTT CAC CGC TTC GAG AAG GAA AAC CGC ATC TGG ACT GTA GTG TTG GAA AGC TAT GTA GTG GAT ATG CCT GAA GGA AAT TCT GAA GAC GAC ACT AGG ATG CTT GCG GAT ACA GTC GTC AAA CTT AAC CTC CAG AAA CTT GCT ACT GTA GCG GAG GCT ATG GCC CGG AAC TCA GGT GAT GGC TCT GGC AGC CAG GTC ACG TGA GGA TCC GCC CCT CTC CCT) by Integrated DNA Technologies, Inc. (IDT, Belgium). Backbone and gBlock were ligated by Gibson assembly and sequenced using standard primers (SV40 for, SV40-pA-rev, IRES rev, EGFP-N-rev).

**TOM20-PYR$^{Mandi}$-IRES-mCherry-ABI** and **TOM20-PYR$^{Mandi}$-IRES-HALO-ABI** were generated from TOM20-mCherry-PYR$^{Mandi}$-IRES-EGFP-ABI. mCherry and EGFP were removed and EGFP was substituted with mCherry or Halo. For TOM20-PYR$^{Mandi}$-IRES-mCherry-ABI, three fragments (backbone, PYR$^{Mandi}$-IRES and mCherry) were amplified from the source vector using flanked primers with appropriate overlaps (#17, #18, #19, #20, #21, #22). For TOM20-PYR$^{Mandi}$-IRES-HALO, the same backbone (#19, #20) and PYR$^{Mandi}$-IRES fragments were used, however PYR$^{Mandi}$-IRES fragment had to be amplified consisting and overlap to Halo fragment (#25, #26). Halo fragment was obtained from TOM20-HALO (#27, #28), which was a gift from Dirk Ollech (Karolinska Institutet). All fragments were ligated by Gibson Assembly and sequenced using standard primers (SV40 for, SV40-pA-rev).

**LCK-CFP-PYL** was generated from LCK-CFP-SNAP$_f$, which was a gift from Dirk Ollech (Karolinska Institutet). SNAP$_f$ was removed by amplification of the backbone (#29, #30). PLY was amplified from TOM20-mCherry-PYL-IRES-EGFP-ABI with appropriate overlaps to the backbone fragments (#31, #32). All fragments were purified by preparative agarose gel and extracted with QIAquick Gel Extraction Kit (Qiagen). All fragments were ligated by Gibson Assembly and sequenced using standard primers (EGFP-C-F-31).

**Vimentin-mNeonGreen-PYR$^{Mandi}$-IRES-Halo-ABI** and **keratin-mNeonGreen-PYR$^{Mandi}$-IRES-Halo-ABI** were cloned from TOM20-PYR$^{Mandi}$-IRES-HALO-ABI. TOM20 side was removed by amplification of the backbone (#33, #34). Fragments of vimentin-mNeonGreen and keratin-mNeonGreen were amplified with Gibson overlaps from mNeonGreen-Keratin-17 (#35, #36) and mNeonGreen-Vimentin-7 (#37, #38), both obtained from Allele Biotech (San Diego, USA). All fragments were ligated by Gibson Assembly and sequenced using standard primers (SV40for).

**TOM20-mCherry-PYL** was cloned from TOM20-mCherry-PYL-IRES-EGFP-ABI. TOM20-mCherry-PLY fragment was amplified by PCR (#39,) with appropriate Gibson overlaps to pcDNA3-swap (pcDNA3 vector with swapped XhoI and XbaI restriction sides, gift from Aline Daniel (Heidelberg University)). pcDNA3-swap was cut with fast digest XhoI and XbaI enzymes (Thermo Fisher Scientific). All fragments were ligated by Gibson Assembly and sequenced using standard primers (T7, SP6).

**TOM20-SNAP$_f$-PYL** was cloned from TOM20-mCherry-PYL. mCherry was removed by amplification of the backbone (#41, #42). SNAP$_f$ fragment was amplified with appropriate Gibson overlaps to the backbone from LCK-CFP-SNAP$_f$, (#43, #44), which was a gift from Dirk Ollech (Karolinska Institutet). All fragments were ligated by Gibson Assembly and sequenced using standard primers (T7).

**AntiGFPNanobody-PYR$^{Mandi}$-IRES-mCherry-ABI** was cloned from TOM20-PYR$^{Mandi}$-IRES-mCherry-ABI. TOM20 sequence was removed by PCR amplification of backbone (#45, #46). AntiGFPNanobody insert was cloned with Gibson overlaps from pOPINE-GFP-nanobody (#47, #48), which was obtained from Addgene (#49172). All fragments were ligated by Gibson Assembly and sequenced using standard primers (SV40-for).

**AntiGFPNanobody-PYR$^{Mandi}$** and **mCherry-ABI** were cloned from AntiGFPNanobody-PYR$^{Mandi}$-IRES-mCherry-ABI by deletion of either IRES-mCherry-ABI (#49, #50) or AntiGFPNanobody-PYR$^{Mandi}$-IRES (#51, #52). Fragments were religated by Gibson Assembly and sequenced using standard primers (SV40-for).

**EGFP-ABI** was cloned from EGFP-GID1 (#37306). GID1 sequence was removed by PCR amplification of backbone (#53, #54). ABI insert with Gibson overlaps was amplified from TOM20-mCherry-PYR$^{Mandi}$-IRES-EGFP-ABI (#55, #56). Fragments were ligated by Gibson Assembly and sequenced using standard primers (EGFP-for).

**TOM20-mCherry-PYR$^{Mandi}$** was cloned from EGFP-GID1 (#37306). EGFP-GID1 sequence was removed by PCR amplification of backbone (#54, #57). PYR$^{Mandi}$ insert with Gibson overlaps was amplified from TOM20-mCherry-PYR$^{Mandi}$-IRES-EGFP-ABI (#58, #59). Fragments were ligated by Gibson Assembly and sequenced using standard primers (EGFP-for).

**LYN-mCherry-PYR$^{Mandi}$** was cloned from TOM20-mCherry-PYR$^{Mandi}$. TOM20 sequence was removed by PCR amplification of backbone (#57, #60). LYN fragment with Gibson overlaps was generated by hybridization of two complementary primers (#61, #62) for 5 min at 95°C. Fragments were ligated by Gibson Assembly and sequenced using standard primers (CMV-for).

**pEGFP-PYL** was cloned from EGFP-ABI. ABI sequence was removed by PCR amplification of backbone (#65, #66). PYL fragment with appropriate Gibson overlaps was amplified from tom20-mCherry-PYL-IRES-EGFP-ABI (#67, #68). Fragments were ligated by Gibson Assembly and sequenced using standard primers (EGFP-for).

**pYFP-PYR$^{Mandi}$** was cloned from LYN-mCherry-PYR$^{Mandi}$. LYN-mCherry sequence was removed by PCR amplification of backbone (#69,#70). YFP fragment with appropriate Gibson overlaps (#71, #72) was amplified from pcDNA3-LAP2-YFP-NLS (#25847). Fragments were ligated by Gibson Assembly and sequenced using standard primers (CMV-for).

### 2 Cell culture, transfection and sample preparation

#### 2.1 General remarks

Cells were grown at 37°C and 5.0% $CO_2$ in Dulbecco's Modified Eagle Medium (DMEM, Sigma-Aldrich, USA) supplemented with 2 mM L-Glutamine, 1 mM Sodium Pyruvate and 10% (v/v) Fetal Bovine Serum. Cells were routinely passaged after 2-3 days or upon reaching 80% confluency.

Prior to seeding cells, type#1 8-well LabTek chambered cover slips (Nunc) were cleaned with 0.1 M hydrofluoric acid to improve attachment of cells. 24 hours before imaging, cells seeded in LabTek chambers were transiently transfected using FuGene HD (Promega, USA) or TransIT-X2 (Mirus, USA) transfection reagents according to manufacturer's guidelines.

Cells transfected with HaloTag or $SNAP_f$-tag fusion constructs were labelled prior to imaging. Growth medium was exchanged with staining solution containing either HaloTag ligand-SiR (HTL-SiR) at 20 nM concentration or TMRStar at 200 nM concentration in DMEM and incubated between 20 min (HaloTag labeling) and up to one hour ($SNAP_f$-tag labeling).

For mitochondria staining, cells were incubated with 200 nM MitoTracker Orange CMTMRos (Thermo Fisher Scientific, USA) in DMEM was incubated for one hour according to manufacturer's guidelines.

All measurements were performed in Leibovitz L-15 medium (Sigma-Aldrich, USA).

All used CIPs were purified by preparative HPLC. Lyophilized products were dissolved in dimethyl sulfoxide. Stocks were diluted in L15 medium. Final DMSO concentrations were kept below 2% for all experiments.

#### 2.2 Protein shuttling and co-localization analysis

Samples were prepared according to general remarks. Imaging was started in 150 µl L15 medium. CIPs were subsequently added as 2x, 3x or 4x stocks in 150 µl L15 medium, respectively. The final DMSO concentration was kept below 2%. Images were acquired after incubation times indicated in the respective figure (Figure 2e, Supplementary Figure 14,15) .

For quantitative analysis of shuttling efficiency, confocal z-stacks were acquired at each time point and individual cells in each stack were segmented manually. Pearson correlation coefficients between receiver channel and both receptor channels were then computed using custom-written ImageJ scripts. First, each image was background-corrected by subtracting a copy of the respective image which was smoothed by convolution with a 20 pixel Gaussian. After background correction, Pearson R was calculated using the ImageJ plugin Coloc2 for each z-slice separately. The final Pearson R for each cell was obtained by averaging R values from individual z-slices.

##### 2.3 Nanobody assisted targeting of chemically induced protein proximity (natCIP)

Stable cell lines Hela (LifeAct-GFP-Halo, gift from Jacob Piehler) and REF (Paxillin-YFP, gift from Ada Cavalcanti-Adam) were transfected with Nanobody-PYR$^{Mandi}$ and mCherry-ABI constructs 24 h prior to imaging. Growth medium was exchanged, and imaging was performed in 200 µl L15 medium. Mandi was added at 2x final concentration in 200 µl L15 medium to a final concentration of 50 nM.

#### 3 Widefield epifluorescence microscopy

Widefield imaging was performed using an inverted epifluorescence microscope equipped with an Apo TIRF 100x NA 1.49 oil immersion objective (both Nikon, Japan). An iChrome MLE-LFA multi laser engine (Toptica Photonics AG, Germany) containing four lasers emitting at 405, 488, 561 and 640 nm was used as the light source and fiber-coupled into the microscope using a TIRF illumination module (Nikon). Focus stabilization in timelapse imaging was achieved using a perfect focus system (PFS3, Nikon). Excitation and emission light were separated using a quad-edge dichroic beamsplitter and emitted light was further filtered using bandpass filters (all AHF Analysetechnik, Germany). Images were acquired using an iXon+ 897 Ultra electron-multiplying CCD camera (Oxford Instruments Andor, United Kingdom) which was also used as timing device to synchronize excitation lasers and camera exposures during imaging with alternating laser excitation. The microscope and all connected devices were controlled using the Micromanager software platform[1]. Typically, images were acquired with 50 ms exposure at 5-10 W/cm$^2$ illumination intensity and at 95 or 146 nm pixel size. Multi-spectral images were acquired using a motorized filter wheel equipped with 525/50 nm (eGFP), 605/70 nm (mCherry, TMR) and 685/70 nm (SiR) bandpass filters.

#### 4 Confocal fluorescence microscopy

Imaging was performed on a commercially available confocal microscope (A1R, Nikon) operated by the Nikon Imaging Center at Heidelberg University. The microscope is built around an inverted, motorized Ti2-E stand, equipped with a galvanometric scanner, a perfect focus system (Nikon) and a stage-top incubation chamber for temperature control and CO2 injection (Tokai Hit, Shizuoka, Japan). A 60x Apo λs NA1.4 oil immersion objective was used for excitation and collection of emitted fluorescence. 488, 561 and 638 nm solid state lasers (all Nikon) were used for excitation, a 405/488/561/640 nm quad band dichroic was used for separating excitation from emission light paths. Typically, 34 µW of 488 nm, 11.5 µW of 561 nm and 195 µW of 638 nm light were used for excitation. Signal from eGFP, mCherry/TMR and SiR was further filtered using 515/30 nm, 595/50 nm and 700/75 nm bandpass filters respectively. For 488 and 561 nm detection channels, GaAsP detectors were used for detection. Detection of signal upon 638 nm excitation was performed using a PMT as detector. A pixel size of 110 nm and scan speeds of 2.4 µsec/pixel with 2x line averaging were applied for all data acquisitions. The pinhole was set to a size of 1.2 Airy units. Z-stacks were recorded with a spacing of 500 nm. Nikon Elements was used to control image acquisition and all connected devices.

### 5 Automated screening of CIP efficiency

*Sample preparation:* COS-7 cells were seeded into 8-well LabTek chambered coverslips. Transient transfection with plasmids expressing both, receiver and receptor domains as cytosolic GFP and TOM20-mCherry fusions linked with an IRES sequence was conducted as described above (appropriate plasmids see table S2). The mCherry-tagged mitochondrial receiver or receptor served as signal for segmentation of mitochondria and to determine area to which cytosolic eGFP-tagged protein was recruited (see below). 24 hours after transfection, cells were washed once with L15 medium and then imaged in L15 at room temperature. CIP solution were freshly prepared from DMSO stocks at 2x final concentration in L15 prior to imaging.

*Acquisition:* Automated timelapse epifluorescence imaging was performed on a Nikon epifluorescence setup described above (see chapter 3. Multi-spectral images were acquired with alternating laser excitation between frames where excitation lasers were controlled by an Arduino microcontroller synchronized to the emCCD camera. Typically, images were acquired with 50 ms exposure per image and variable lag times between individual image pairs depending on the typical times-to-effect for the individual CIP systems (Table S2). The end point of the timelapse was chosen so that no further recruitment of the cytosolic signal to mitochondria was observed. The total number of images per post-CIP addition timelapse was kept constant to minimize differences due to photobleaching or phototoxicity between CIPs. eGFP and mCherry were excited with CW laser illumination at 0.35 and 0.51 mW output at the objective corresponding to an average irradiance of 5.3 and 8.6 W/cm$^2$ across the read-out region. Emitted fluorescence was split using an Optosplit II image splitter (Cairn Research, United Kingdom) equipped with a 560 nm longpass beamsplitter (AHF Analysetechnik, Germany) and additionally filtered using 605/70 nm (mCherry) and 525/50 nm (eGFP) bandpass filters inserted in the reflected and transmitted light paths respectively. Signals from both paths were recombined using a second 560 nm shortpass filter and two two-axis translation mirrors. Manual coarse alignment of both channels was achieved using 0.1 µm Tetraspek mutli-fluorescent beads (Thermo Fisher, USA) as reference. For each experiment, a single cell in each well was manually selected with receiver and receptor expression and general cell morphology as selection criteria. Automated data acquisition was then performed using a custom-written µManager[1] beanshell script. In brief, each acquisition consists of a 488/561 nm excitation image pair before CIP addition ($t_0$), CIP addition, timelapse acquisition and acquisition of a final $t_{end}$ image pair (see Supplementary Fig 4a). Injection of CIP was performed with a computer-controlled Alladin AL1000 microfluidic pump (World Precision Instruments, USA) at a flow rate of 6 ml/min. CIP was added at equal volume and double final concentration followed by a 2-4 sec delay to allow for mixing of medium in well and added CIP solution. This procedure was repeated for each well on one slide.

*Data pre-processing:* Acquired multi-dimensional image stacks were processed in Fiji[2] using custom-written analysis routines. Raw data was automatically checked for errors in illumination sequences and corresponding image pairs were removed from timelapse datasets. Across all acquisitions, <1 % of image pairs were discarded during this step. Flat fielding to correct for differences in excitation intensity was performed by multiplying all images with a template image. Illumination profile templates were obtained by acquiring 20-30 images of surfaces homogeneously coated with Alexa Fluor 488 or tetramethyl rhodamine (TMR) NHS esters for 488 and 561 nm excitation respectively. Images were then averaged and normalized to the maximum value in the averaged image. To correct for variations in alignment of the microscope, new templates were measured for each round of experiments. Image pairs were spatially aligned with sub-pixel accuracy using the Image Stabilizer Plugin authored by Kang Li (http://www.cs.cmu.edu/~kangli/code/Image_Stabilizer.html). 0.1 µm Tetraspek beads served as reference structure to compute transformation coefficients. Transformation coefficients were determined separately for each experiment.

*Segmentation & intensity extraction:* Raw image data was automatically segmented using the Trainable Weka Segmentation package[3]. Models for classification of total cell area and mitochondria were trained by manual classification of 10 randomly selected images from the entire dataset. The model for cell detection was trained and applied using 488 nm excitation $t_0$ images which exhibit purely cytosolic signal. The mitochondria model was trained using 561 nm excitation $t_0$ images. The obtained segmentations were robust with respect to the average area occupied by mitochondria in any given image set which typically was between 10 and 40% (Supplementary Fig. 4b) and no systematic variation in segmented mitochondrial area in individual timelapse datasets was observed (Supplementary Fig. 4c).

Regions of interest (ROIs) were obtained from Weka segmentation results by thresholding of segmentation maps. Cytosolic ROIs were obtained by computing the difference between whole cell and mitochondria for each image pair. Since the cytosolic signal gradually translocated to mitochondria after CIP addition, cytosolic ROIs for timelapse and $t_{end}$ image pairs were computed using the $t_0$ whole cell ROI and the mitochondria from current image pair. As expected, the whole cell intensity in the 488 nm channel remained unchanged (<2% variation) after CIP addition indicating that cell movement during timelapse image acquisition was negligible (Supplementary Fig. 4d). After segmentation, average intensities for each ROI and image were extracted and exported as text files. Acquisition time stamps were extracted from image meta data and included in text files.

*Plotting:* Translocation ratios were computed using intensities extracted from $t_0$, $t_{end}$ image pairs and timelapse data. All calculations and plots were created using Matlab 2018a (The MathWorks, USA). In afirst step, all data was corrected for photobleaching using the decay in whole frame intensity during acquisition. Then, ratio of mean 488 nm intensity in the mitochondria and cytosol ROI for each frame of the timelapse dataset was computed. The obtained ratios were corrected for the ratio before CIP addition ($ratio_{t0}$) and the final ratio after timelapse acquisition ($ratio_{tend}$). No ratio was computed for frames with erroneous illumination sequence. Datasets where segmentation was not reliable were identified using the average mitochondrial 561 nm signal over time and excluded from analysis. The fraction of cells excluded was <10% across all CIPs (Supplementary Table 2).

All code required for acquisition and processing of raw data will be made available after final revision of manuscript.

**Supplementary Table 2: Experimental design for automated measurements of CIP-induced protein translocation.**

| CIP | CIP concentration [µM] | Acquisition frame rate [pair/min] | Image pairs per acquisition | Plasmid | Fraction cells successfully processed [%] |
|---|---|---|---|---|---|
| $GA_3$-AM | 5 | 12 | 120 | eGFP-GID1-IRES-TOM20-mCherry-GAI | 96.8 |
| ABA | 5 | 8 | 120 | TOM20-mCherry-PYL-IRES-eGFP-ABI | 92.0 |
| ABA-AM | 5 | 12 | 120 | TOM20-mCherry-PYL-IRES-eGFP-ABI | 93.8 |
| Mandi | 5 | 24 | 120 | TOM20-mCherry-$PYR^{Mandi}$-IRES-eGFP-ABI | 91.7 |
| Mandi | 0.5 | 60 | 300 | TOM20-mCherry-$PYR^{Mandi}$-IRES-eGFP-ABI | 100 |
| Mandi | 0.05 | 60 | 300 | TOM20-mCherry-$PYR^{Mandi}$-IRES-eGFP-ABI | 100 |

### 6 Raster spectral image correlation spectroscopy (RSICS)

*Sample preparation:* For RSICS experiments, $10^5$ COS-7 cells were seeded in 35 mm #1.5 optical glass bottom dishes (CellVis, Mountain View, CA) 24 h before transfection. Cells were co-transfected with 10 ng YFP-PyrMandi, 200 ng eGFP-Pyl, 250 ng mCherry-ABI and imaged 20 h after transfection. For the negative cross-correlation control, cells were co-transfected with 50 ng of mEGFP-N1, YFP-N1 and mCherry-N1 vectors. To calibrate the maximum cross-correlation of the setup, positive control samples were prepared by transfecting cells with 50 ng of mCherry-eGFP or mCherry-YFP heterodimer constructs, as described previously[4]. For single species samples, cells were transfected with 50 ng of either mEGFP-N1, YFP-N1 or mCherry-N1. All transfections were performed using Lipofectamin3000 according to the manufacturer's instructions (Thermo Fisher Scientific). Further information on plasmids for control measurements can be found in Dunsing *et al.*[4]. RSICS measurements with ABA-AM and Mandi were performed after 15 min incubation of samples supplemented with 5 µM ABA-AM or Mandi.

*Data acquisition:* RSICS measurements were performed on a Zeiss LSM880 system (Carl Zeiss, Oberkochen, Germany) using a 40x, 1.2NA water immersion objective. Per measurement, 300-400 frames of 256x256 pixels were acquired with 50 nm pixel size (i.e. a scan area of 12.83x12.83 µm through the midplane of cells), 2.05 µs pixel dwell time, 1.23 ms line and 314.57 ms frame time (corresponding to ca. 1.5-2 min total acquisition time). Samples were excited with a 488 nm Argon laser and a 561 nm diode laser at ca. 4.8 µW (488 nm) and 5.9 µW (561 nm) excitation powers, respectively. Laser powers were chosen to maximize the signal emitted by each fluorophore species but keeping photobleaching below 25% for all species. Typical counts per molecule were ca. 25 kHz for eGFP (G), 15-20 kHz for YFP (Y) and 10 kHz for mCherry (Ch). To split excitation and emission light, a 488/561 nm dichroic mirror was used. Fluorescence was detected between 490 nm and 695 nm in 23 spectral channels of 8.9 nm on a 32 channel GaAsP array detector operating in photon counting mode. To obtain reference emission spectra for each individual fluorophore species, 4 image stacks of 25 frames were acquired at the same imaging settings on single species samples on each day. In addition, negative and positive cross-correlation controls samples were measured on each day. All measurements were performed at room temperature.

*Data analysis:* RSICS analysis followed the implementation described recently[5], which is based on applying the mathematical framework of fluorescence lifetime and fluorescence spectral correlation spectroscopy[6,7] to raster image correlation spectroscopy (RICS). Four-dimensional image stacks were imported in MATLAB (The MathWorks, Natick, MA) from CZI image files using the Bioformats package[8] and further analyzed using custom-written code. First, average reference emission spectra were calculated for each individual fluorophore species from single species measurements. Four-dimensional image stacks were then decomposed into three three-dimensional image stacks (G, Y, Ch) using the spectral filtering algorithm presented by Schrimpf *et al.*[5]. Cross-correlation RICS analysis was performed in the arbitrary region RICS (ARICS) framework[9]. To this aim, a region of interest was selected in the time- and channel-averaged image frame containing a homogeneous region in the cytoplasm of cells. This approach allowed excluding visible intracellular organelles or pixels in the extracellular space. Image stacks were further processed with a high-pass filter (with a moving 4-frame window) to remove slow signal variations and spatial inhomogeneties. Afterwards, RICS autocorrelation functions (ACFs) and three pair-wise cross-correlation functions (CCFs) were calculated for each image stack and the three detection channel combinations G-Y, G-Ch, Y-Ch, respectively[5,9]. A normal diffusion RICS fit model[10,11] was then fitted to both, ACFs and CCFs. From the amplitudes of the ACFs and CCFs, the relative cross-correlation was calculated for the three cross-correlation (CC) combinations G-Y, G-Ch, Y-Ch:

$$rel.cc. = max\left\{\frac{G_{CC,ij}(0)}{G_{AC,i}(0)}, \frac{G_{CC,ij}(0)}{G_{AC,j}(0)}\right\},$$

where $G_{CC,ij}(0)$ is the amplitude of the CCF of species *i* and *j*, and $G_{AC,i}(0)$ the amplitude of the ACF of species *i*.

Binding efficiencies were calculated by subtracting the residual average relative cross-correlation measured in each cross-correlation channel of the negative control (containing three mixed FP species) from the measured cross-correlation. The result was then normalized using the average relative cross-correlation obtained from the positive cross-correlation controls (containing eGFP-mCherry or eGFP-YFP heterodimers).The positive controls account for imperfect alignment of the optical observation volumes as well as non-fluorescent states of the fluorescent protein tags (*e.g.* due to limited maturation or dark states[4,12]). To ensure statistical robustness of the three-color RICS analysis and sufficient signal-to-noise ratios, the analysis was restricted to cells expressing all three fluorophore species in comparable amounts, *i.e.* relative average signal intensities of less than 3 for all species.

All code required for processing of raw data can be made available upon reasonable request after final revision of manuscript.

### 7 In vivo Experiments

#### 7.1 Zebrafish strains

The $AB_2O_2$ WT line (European Zebrafish Resource Centre EZRC, Karlsruhe) was used for all experiments. Zebrafish husbandry[13] and experimental procedures were performed in accordance with German animal protection regulations (Regierungspräsidium Karlsruhe, Germany, 35-9185.64/BH KIT).

#### 7.2 Real-time imaging of zebrafish embryos

EGFP-ABI and TOM20-mCherry-$PYR^{Mandi}$ or LYN-mCherry-$PYR^{Mandi}$ plasmids were injected into the yolk of 1–2 cell embryos[14]. Positive co-expressing 3- to 5-day-old embryos, which were immobilized on a microscopy slide using 0.5% low melting point agarose supplemented with 0.02% MESAB were used. Embryos were imaged with a water dip-in 63x objective (NA: 0.90; HCX APO water; Leica) and installed at a Leica TCS SP2 confocal microscope and the corresponding Leica LCS software. All experiments were performed at room temperature. Mandi in water (50 µM stock solution in 100% DMSO; 500 nm final concentration) was added on top of the embedded embryos.

### 8 Supplementary Figures

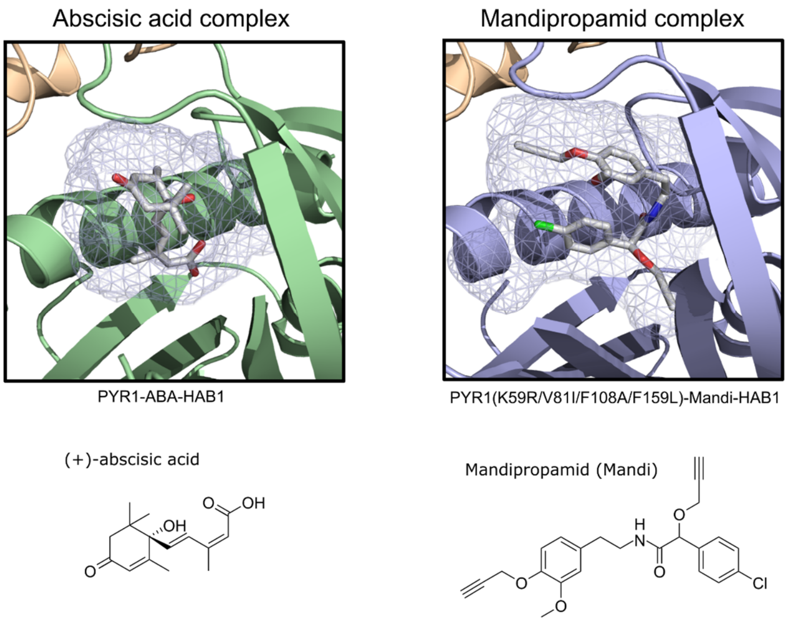


**Supplementary Figure 1: Comparison of crystal structures of abscisic and mandipropamid binding sites in receptor and receiver complex and chemical structures of CIP[15,16].** Binding receptor for Mandi quadruple mutant PYR(K59R/V81I/F108A/F159L) as for sextuple mutant PYR$^{Mandi}$ no crystal structure is reported. GEHRL-loop in front of binding pocket has been omitted for better visualization. Crystal structures were obtained from pdb entries 3JRQ and 4WVO.

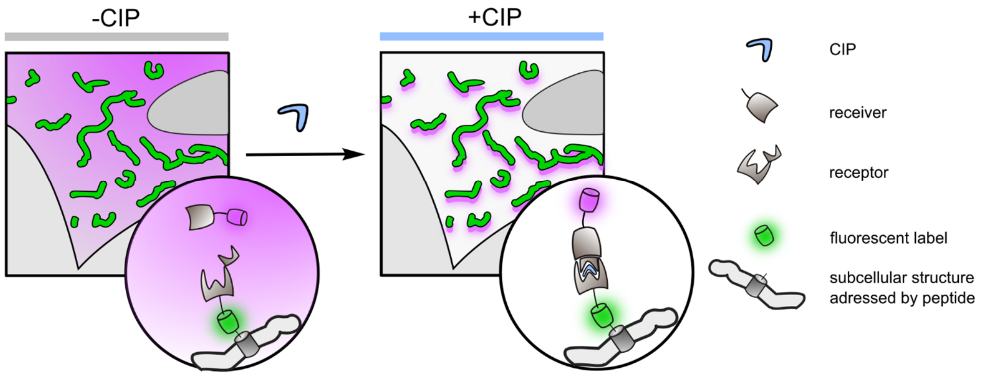


**Supplementary Figure 2: Schematic of colocalization assay.** Receptor or receiver is expressed as fusion to proteins with known localization. Protein interaction is under strict control of CIP addition. Fluorescent labels (fluorescent proteins or organic dyes introduced by protein tag strategies) allow monitoring colocalization by fluorescence microscopy.

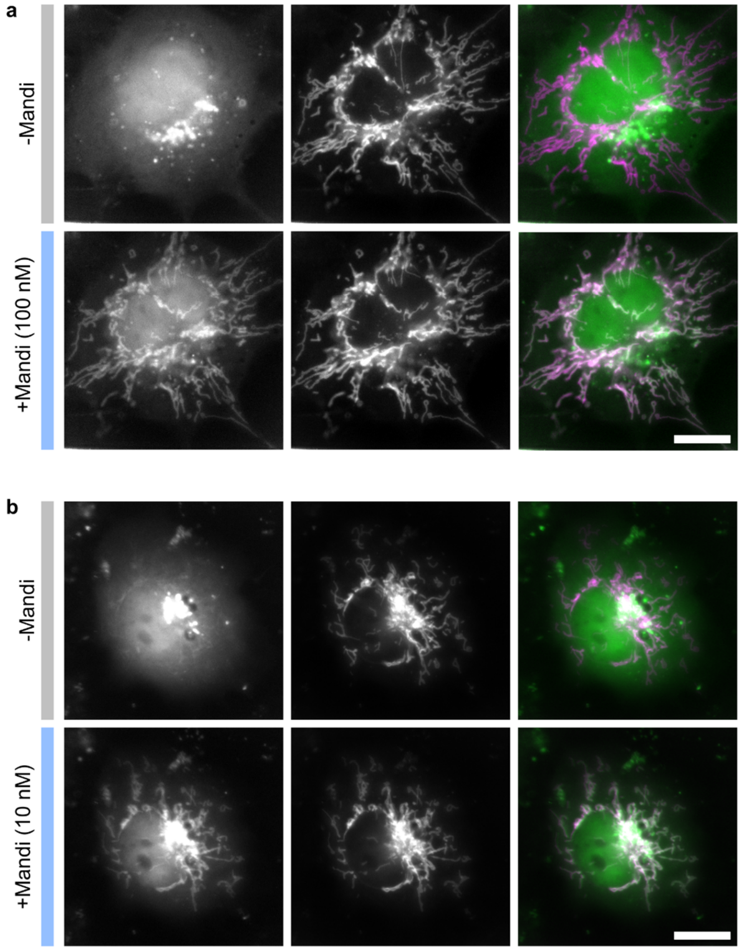


**Supplementary Figure 3: Live-cell epifluorescence microscopy images of Mandi-induced colocalization using different concentrations.** COS-7 cells were transfected with TOM20-mCherry-PYR$^{Mandi}$-IRES-EGFP-ABI. Images were acquired before addition of 100 nM (a) and 10 nM (b) Mandi and after completion. Timelapse of colocalization process shown in supplementary video 1,2. Colocalization was completed within 1 min (a) and 3-5 min (b). Scale bar 20 µm. Representative data for a) n=6 cells b) 12 cells from 2 independent experiments.

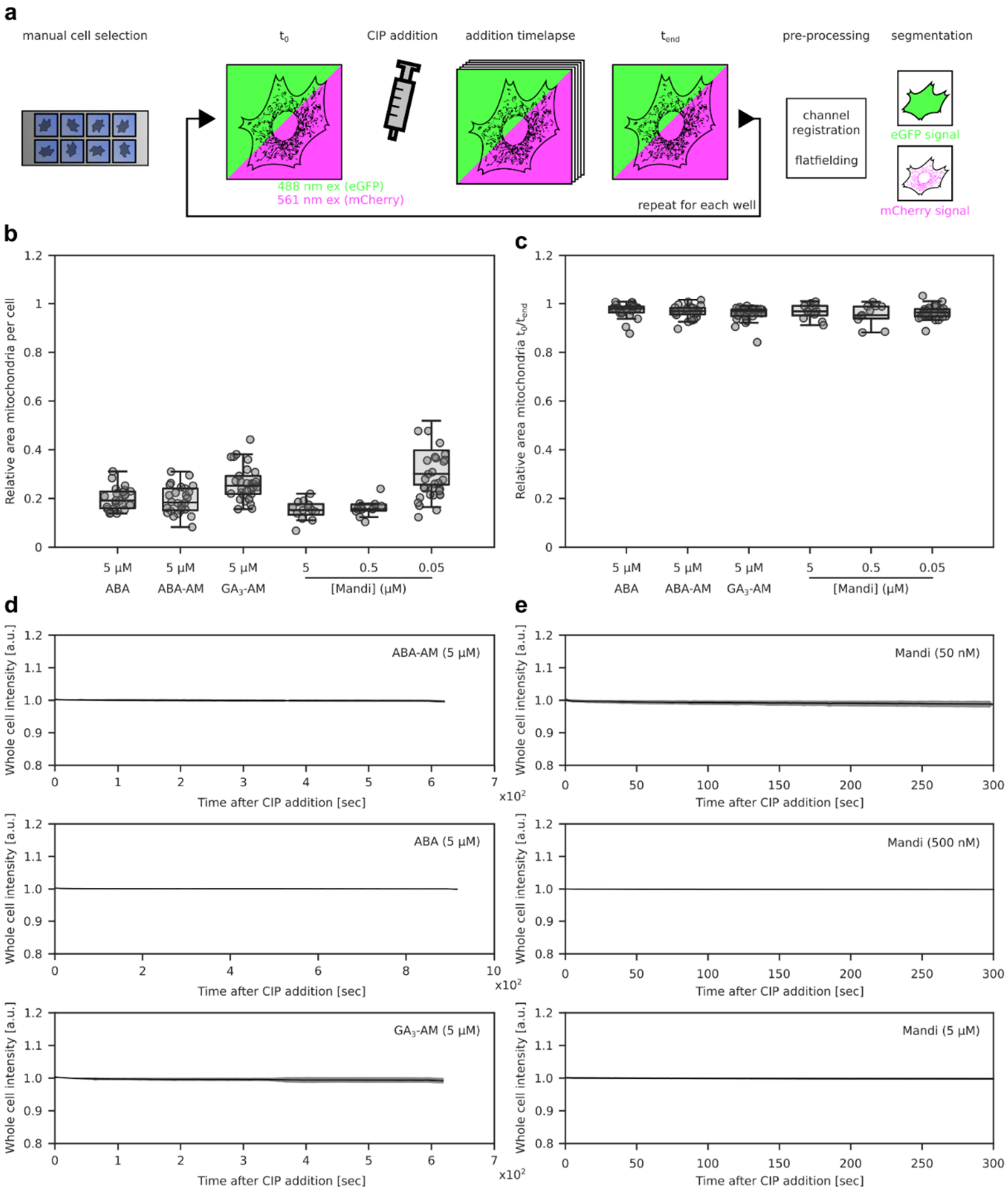


**Supplementary Figure 4: Screening of CIP efficiency with automated microscopy. a,** experimental workflow consisting of manual cell selection, automated microscopy and CIP addition and data processing. **b,** relative area occupied by mitochondria obtained from calculating the fraction of cell ROI vs. mitochondria ROI. Box plot indicates 25th percentile, median and 75$^{th}$ percentile. Whiskers extend to 1x interquartile distance. **c,** relative mitochondrial ROI size in $t_0$ and $t_{end}$ image pairs obtained from thresholding Weka probability maps. **d,** normalized change in 488 nm intensity measured across cell ROI for all cells with a given CIP. Solid line indicates average and shaded region variation (±1 standard deviation) over time. Data in (b,c,d) from datasets shown in **Fig. 1g,h**.

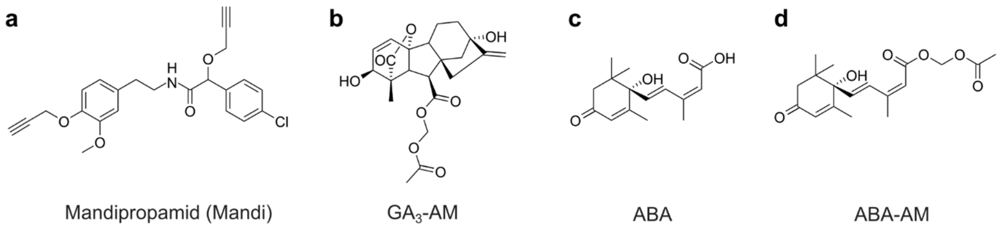


**Supplementary Figure 5: Chemical structures of different CIPs used in CIP screen. a,** Mandipropamid (Mandi), **b,** Gibberellic acid acetoxymethyl ester ($GA_3$-AM), **c,** abscisic acid (ABA), **d,** abscisic acid acetoxymethyl ester (ABA-AM).

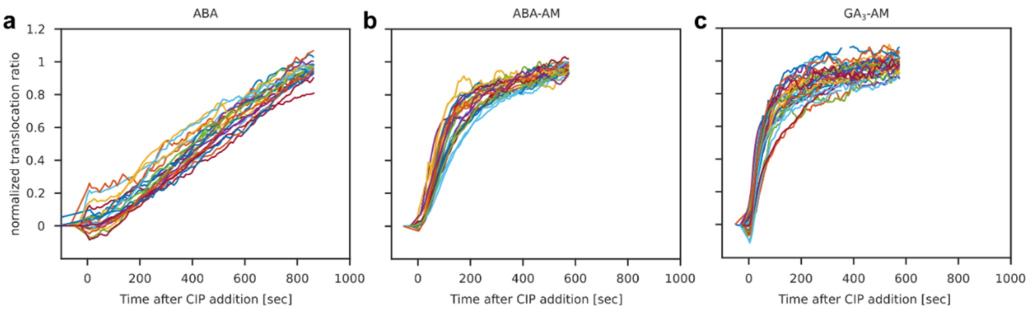


**Supplementary Figure 6: Single-cell translocation ratios. a,** ABA. **b,** ABA-AM. **c,** $GA_3$-AM. Each line represents photobleaching-corrected intensity ratio from mitochondria and cytosol normalized by corresponding $t_0$ and $t_{end}$ values. Gaps in individual traces due to discarded frames (see methods). CIP addition at time t=0 sec.

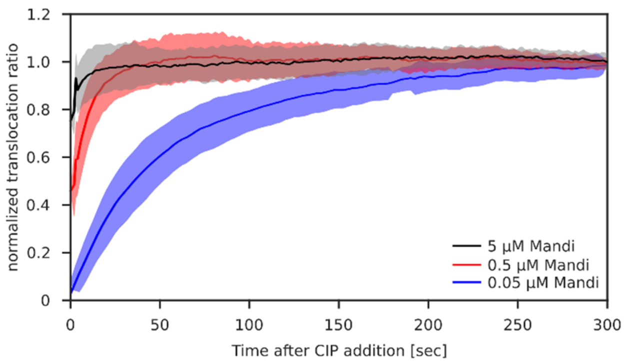


**Supplementary Figure 7: Averaged translocation ratios for different Mandi concentrations.** Source data used to compute translocation times $t_{0.75}$ shown in **Fig 1h**. Solid line indicates average and shaded region variation (±1 standard deviation) over time.

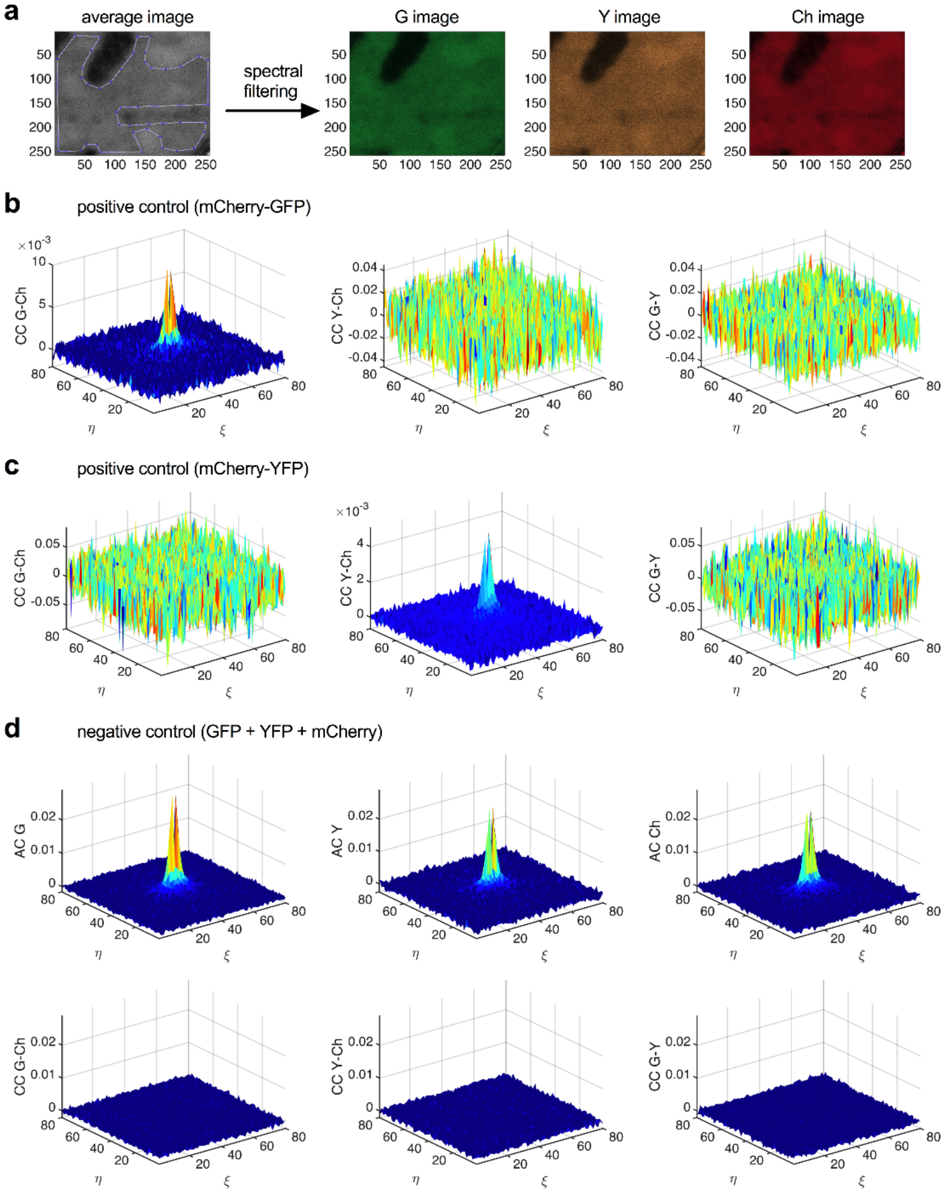


**Supplementary Figure 8: Three-color RSICS measurements of positive and negative cross-correlation controls. a,** Schematic of RSICS in the ARICS framework. A 256x256 pixel image stack of 300-400 frames acquired in 23 spectral channels is decomposed into three three-dimensional image stacks for the three species GFP (G), YFP (Y) and mCherry (Ch), using a spectral filtering algorithm. An arbitrary ROI delimiting a homogeneous region in the cytoplasm is selected and RSICS analysis is applied to each frame of the three image stacks in the ARICS framework. **b,c** RSICS CCFs in the three cross-correlation channels G-Ch, Y-Ch and G-Y measured on COS-7 cells expressing fluorescent constructs used as positive controls for intermolecular interactions: mCherry-GFP **(b)** and YFP-mCherry **(c)** hetero-dimers. **d,** ACFs (top row) and CCFs (bottom row) obtained from RSICS measurement on COS-7 cell co-expressing fluorescent constructs used as negative controls for inter-molecular interactions, *i.e.* free non-interacting GFP, YFP, and mCherry fluorophores.

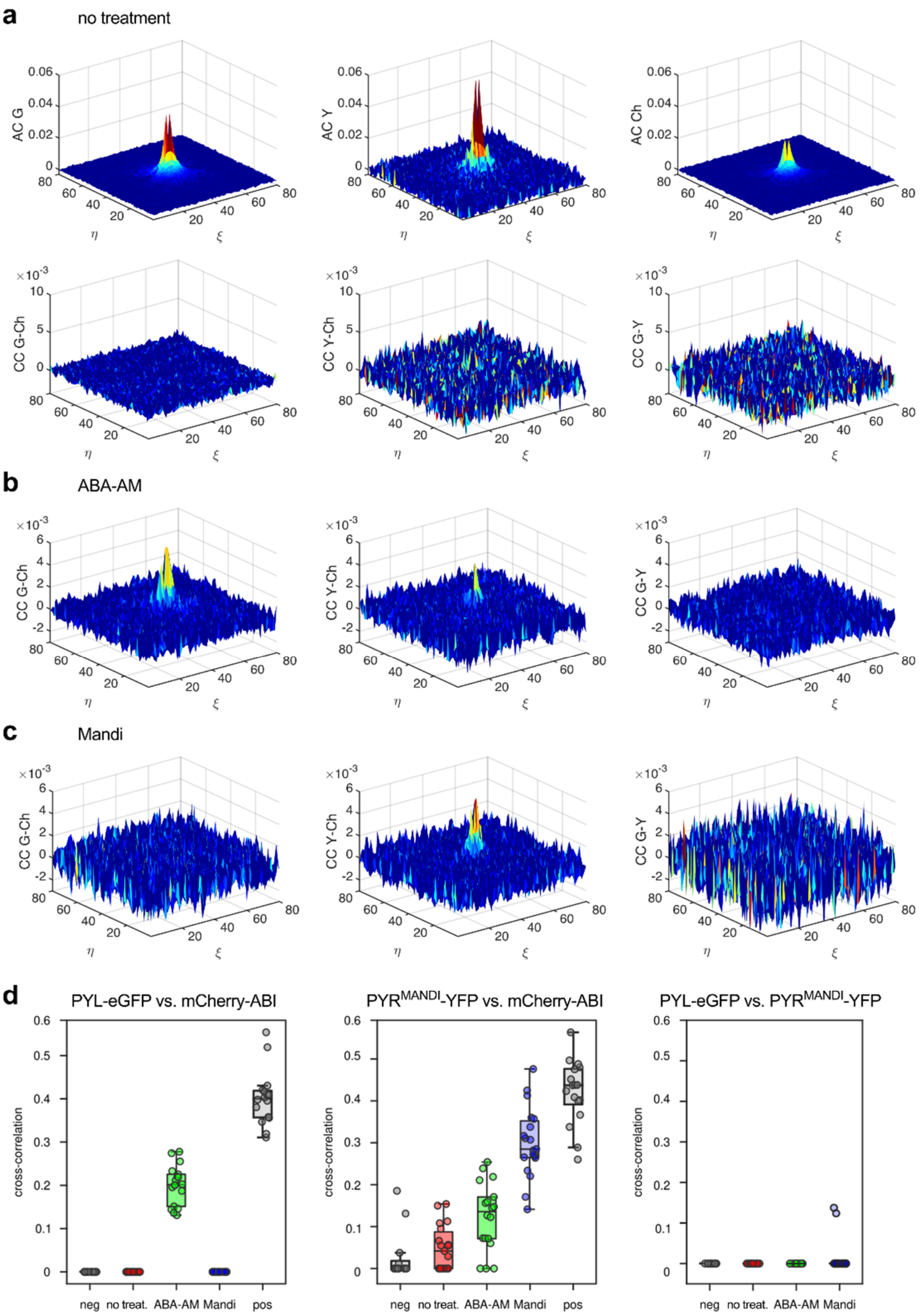


**Supplementary Figure 9: Three-color RSICS measurements on the Mandi CIPP system. a,** ACFs (top row) and CCFs (bottom row) obtained from RSICS measurement on COS-7 cells co-expressing PYL-GFP, $PYR^{MANDI}$-YFP and mCherry-ABI before treatment with ABA-AM or Mandi. **b,c** CCFs in the three cross-correlation channels obtained from RSICS measurement on COS-7 cells co-expressing PYL-GFP, $PYR^{MANDI}$-YFP and mCherry-ABI performed 15 min after incubation with 5 µM ABA-AM **(b)** or 5 µM Mandi **(c)**. **d,** Box plots of relative cross-correlation values measured in the three cross-correlation channels pooled from all RSICS measurements on the Mandi CIP system (*i.e.* G-Ch/ Y-Ch/ G-Y, corresponding to interactions between PYL-GFP and mCherry-ABI/ PYR$^{Mandi}$-YFP and mCherry-ABI/ PYL-GFP and PYR$^{Mandi}$-YFP) in the absence (no treat.) and presence of 5 µM ABA-AM or 5 µM Mandi. For comparison, the cross-correlation values obtained in the negative (neg) and positive (pos) cross-correlation control samples are shown. Each data point corresponds to one RSICS measurement in a single cell. Data are pooled from two independent experiments with 13 (neg), 19 (no treat.), 18 (ABA-AM), 19 (Mandi) and 16 (pos) cells.

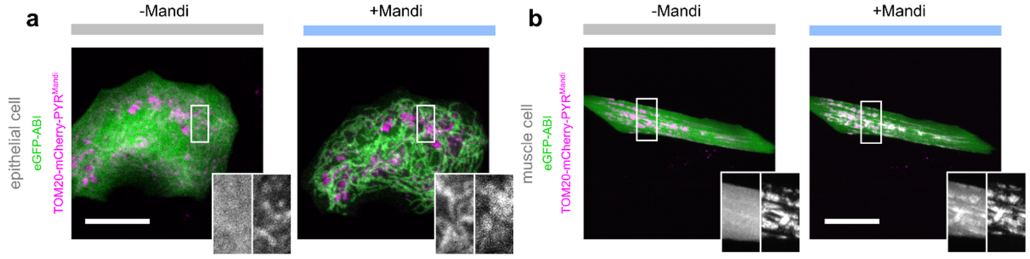


**Supplementary Figure 10: Confocal fluorescence microscopy images of protein colocalization on different cell types in living zebrafish embryos.** Receiver and mitochondria localized receptor domains expressed in zebrafish embryos. **a,** epithelial cell **b,** muscle cell. Images acquired before and 10-20 min. after addition of 500 nM Mandi. Scale bar 40 µm. Data representative for n=3 independent experiments.

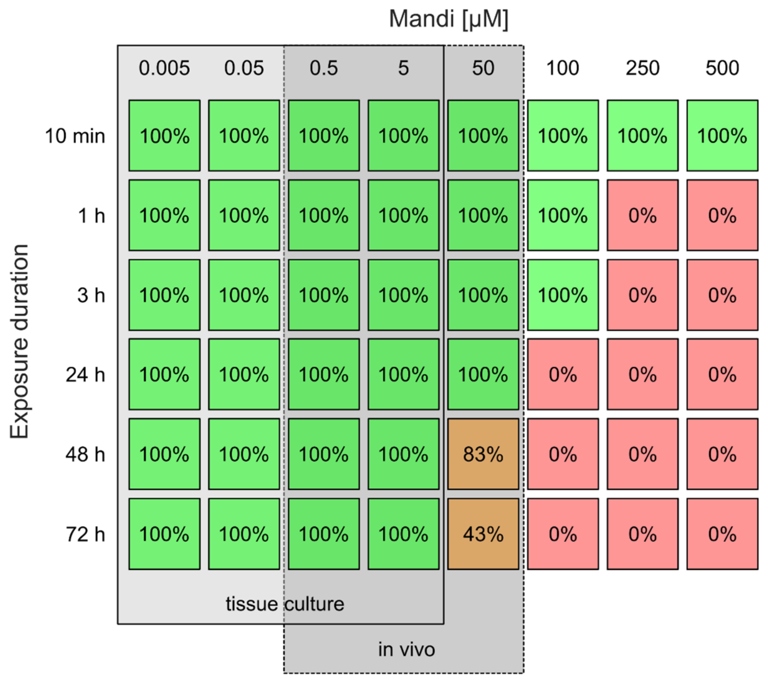


**Supplementary Figure 11: Toxicological investigation of Mandi influence on zebrafish embryos.** Investigation started 3-5 dpf. Survival rate summarized from 3 independent experiments using 10 embryos per condition. High concentrations of 500 µM to 50 mM showed phenotypes such as accumulation of red blood cells in the heart and slowed heartbeat.

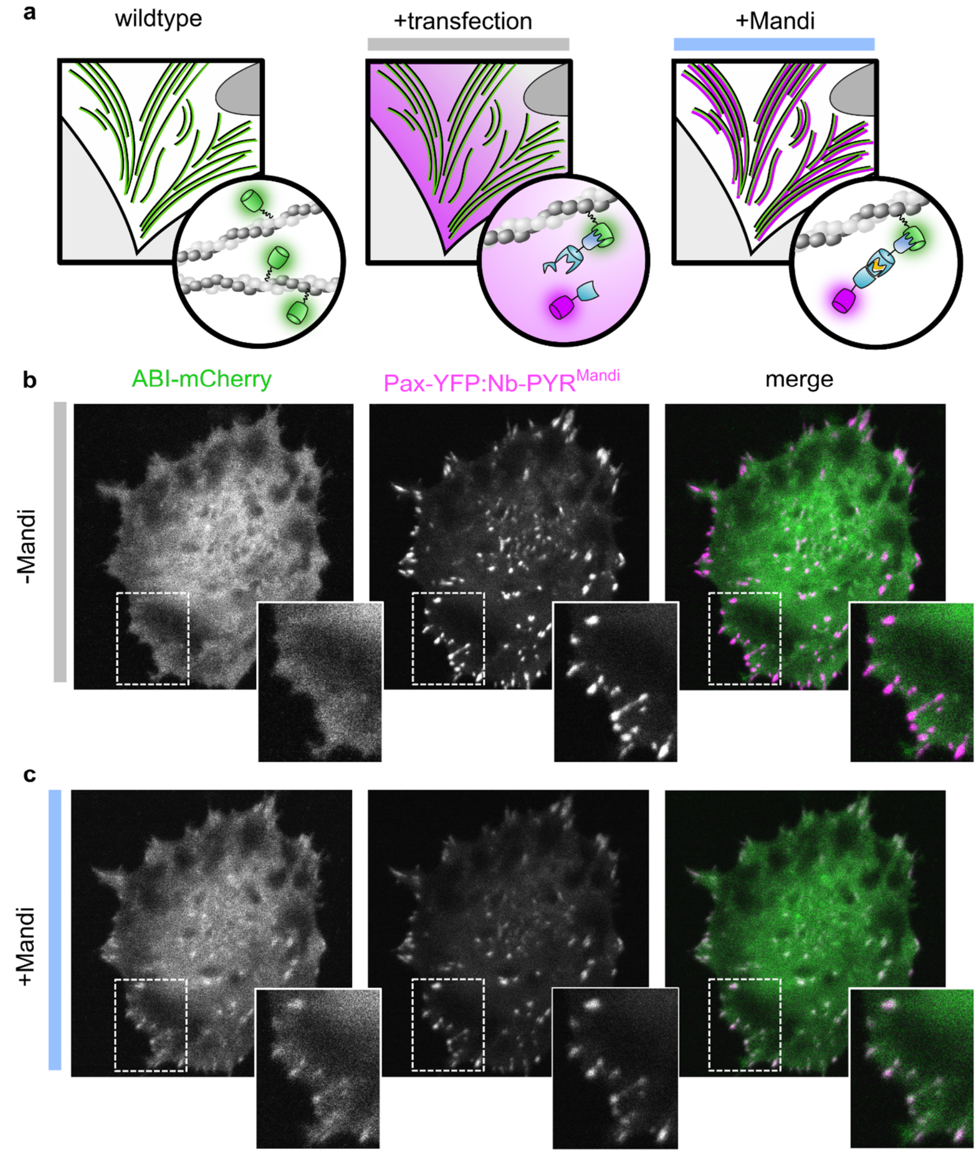


**Supplementary Figure 8**: **natCIPP in rat embryonic fibroblast (REF) cells stably expressing paxillin-YFP. a,** Schematic illustration of natCIPP. REF cells stably expressing paxillin-GFP were transfected with antiGFP-nanobody-PYR$^{Mandi}$ and mCherry-ABI fusions. **b,** Confocal fluorescence microscopy images acquired before and **c,** 5 min after addition of 50 nM Mandi. Scale bar 20 µm. Representative data for n=15 cells from 2 experiments.

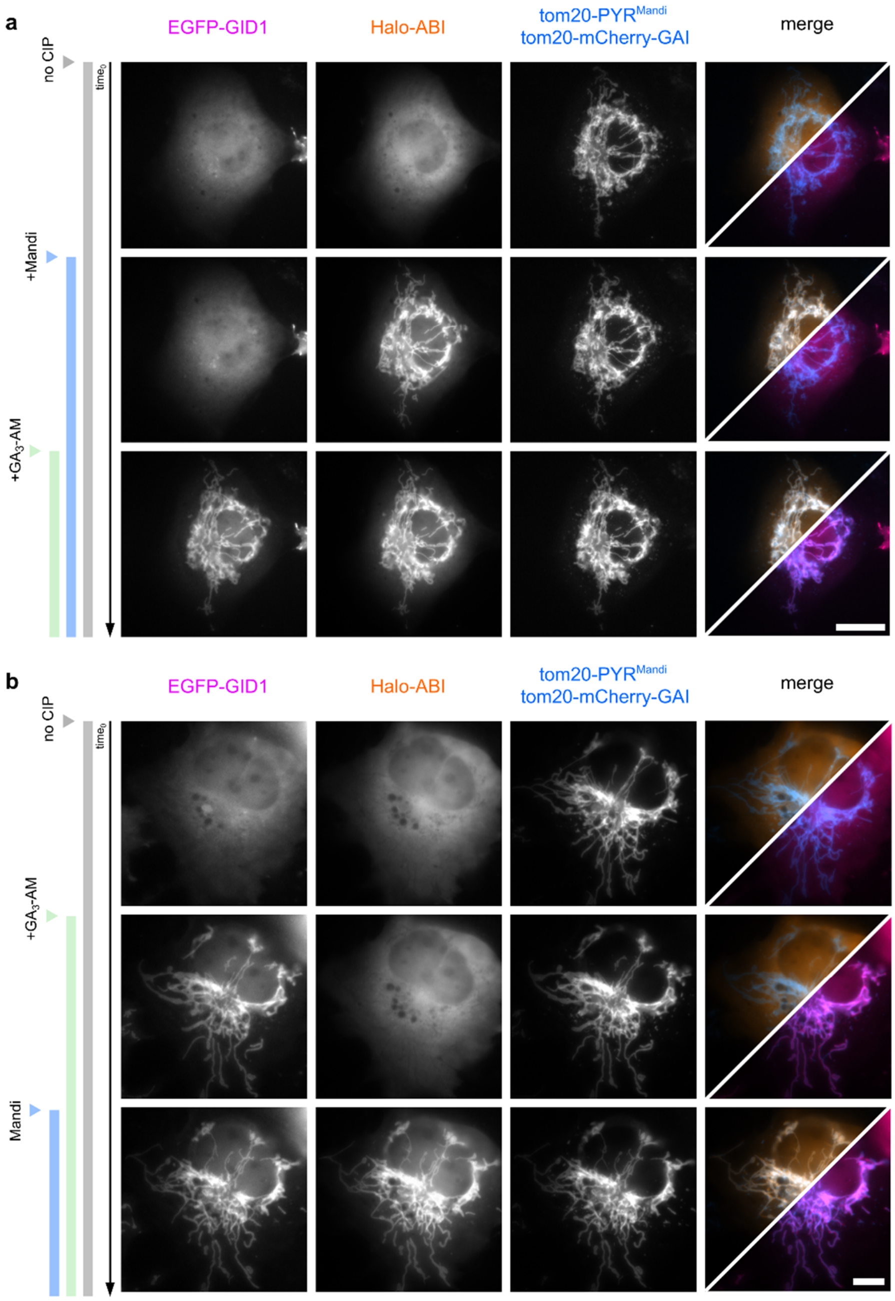


**Supplementary Figure 9: Live-cell epifluorescence microscopy images to show orthogonality of Mandi and $GA_3$-based CIPP systems.** COS-7 cells were co-transfected with TOM20-mCherry-GAI-IRES-EGFP-GID1 and TOM20-PYR$^{Mandi}$-IRES-Halo-ABI. Halo-tag domain was stained with HTL-SiR prior to imaging. **a,** Images acquired before CIP addition (row 1), 5 minutes after addition of Mandi (50 nM, row 2), 5 min after addition of $GA_3$-AM (500 nM, row 3). **b,** Images acquired before CIP addition (row 1), 5 minutes after addition of $GA_3$-AM (500 nM, row 2), 5 min after addition of Mandi (50nM, row 3). Scale bar 10 µm. Representative data for n=3 cells.

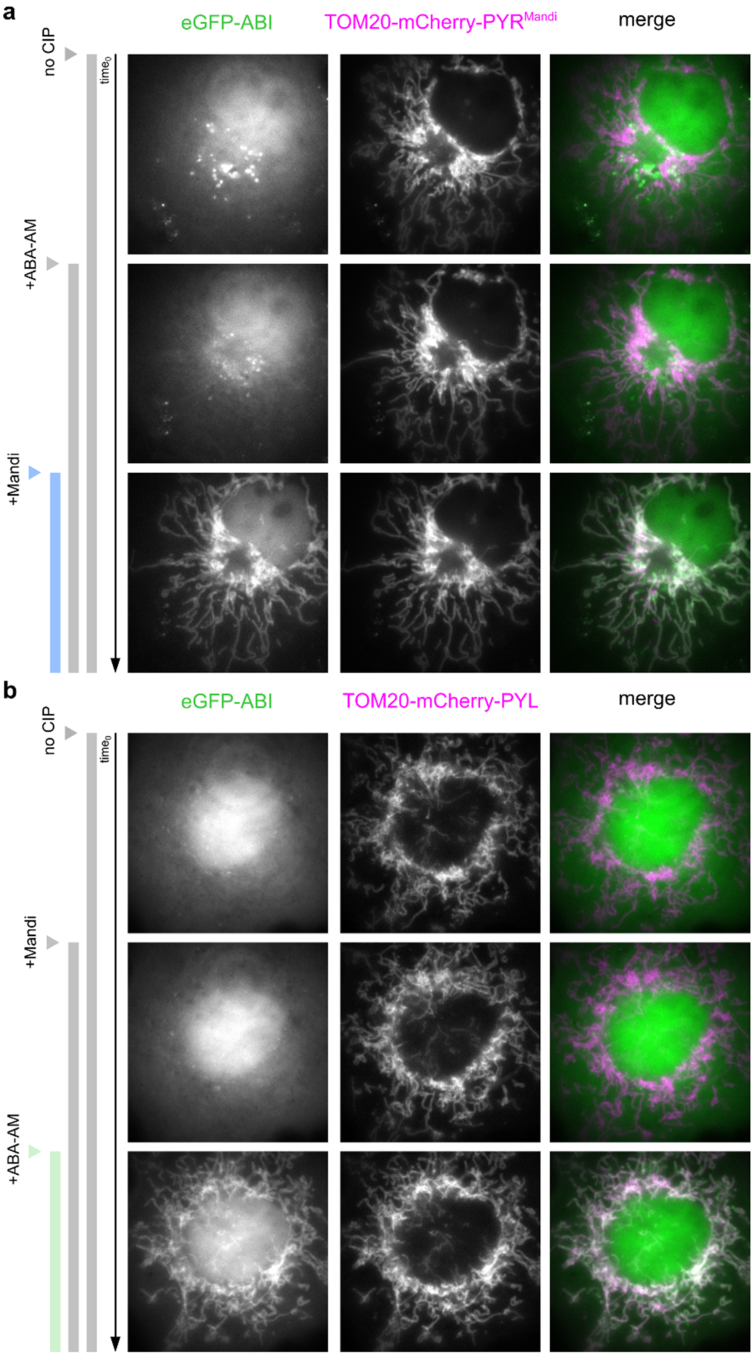


**Supplementary Figure 10: Live-cell epifluorescence microscopy images to show semi-orthogonality of Mandi and ABA- based CIPP systems. a,** COS-7 cells were transfected with TOM20-mCherry-PYR$^{Mandi}$-IRES-EGFP-ABI. Images acquired before CIP addition (row 1), 10 minutes after addition of ABA-AM (200 nM, row 2), 10 min after addition of Mandi (200 nM, row 3). **b,** COS-7 cells were transfected with TOM20-mCherry-PYL-IRES-EGFP-ABI. Images acquired before CIP addition (row 1), 10 minutes after addition of Mandi (200 nM, row 2), 10 min after addition of ABA-AM (200 nM, row 3). Scale bar 10 µm. Representative data for n=11 cells**.**

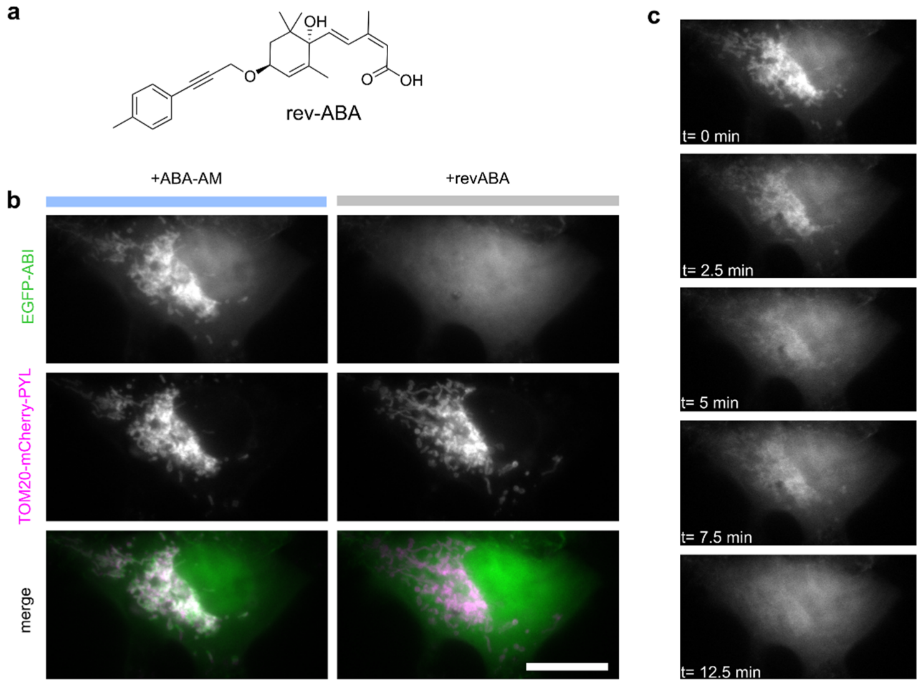


**Supplementary Figure 11: revABA can reverse ABA-AM induced protein-protein interaction. a,** Molecular structure of revABA. **b,** Epifluorescence microscopy images of COS-7 cells transfected with TOM20-mCherry-PYL-IRES-EGFP-ABI. Cells were incubated for 4 h with ABA-AM (200 nM) to induce dimerization. Imaging was performed in L15 with ABA-AM (200 nM). Images were acquired before and 20 min after addition of revABA (5 µM, 50x excess). **c,** Exemplary images from timelapse (Supplemenary video 4). Reversion completed after ~12.5 min. Scale bar 20 µm. Representative data for n=20 cells from 3 experiments.

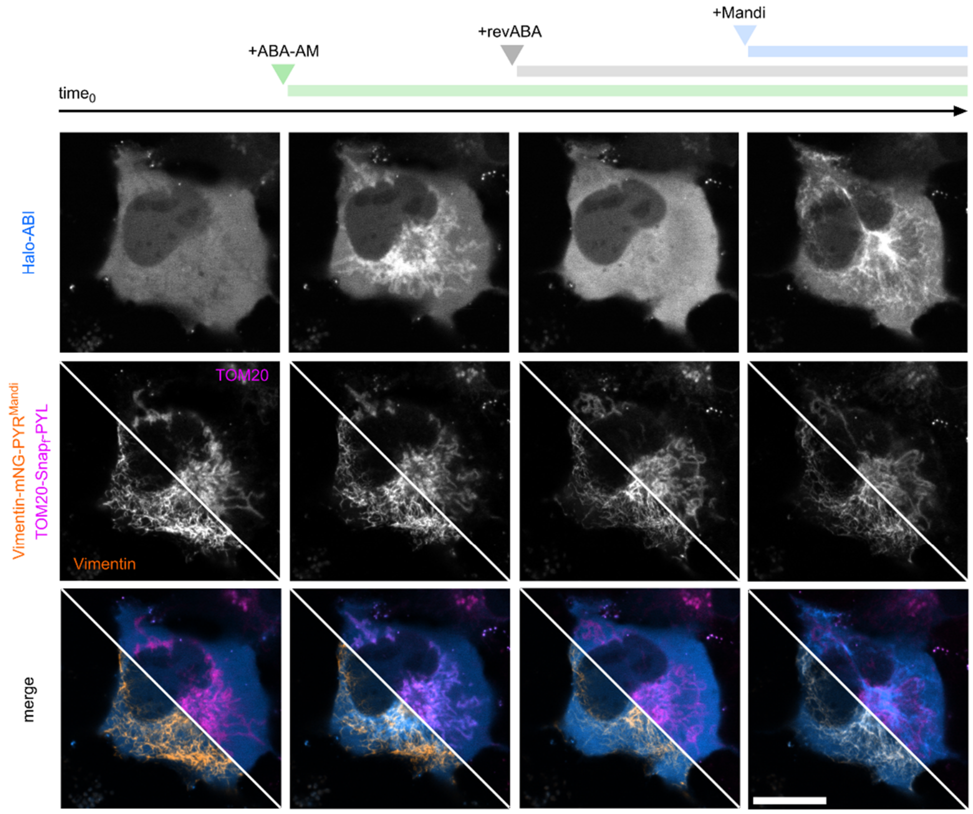


**Supplementary Figure 12: Reversible and dynamic protein shuttling between mitochondria and vimentin in living cells.** COS-7 cells were co-transfected with Vimentin-mNeonGreen-PYR$^{Mandi}$-IRES-Halo-ABI and TOM20-SNAP$_{f}$-PYL. Halo- and SNAP-tag domains were stained with HTL-SiR and TMRStar 2 h prior to imaging. Upper row shows dynamic receiver localization, middle row receptor localizations as references, lower row respective merges. Images acquired at $t_0$, 10 min after addition of ABA-AM (200 nM), 25 min after addition of revABA (20 µM), 10 min after addition of Mandi (200 nM). Scale bar 20 µm. Representative data for n=22 cells from 2 independent experiments.

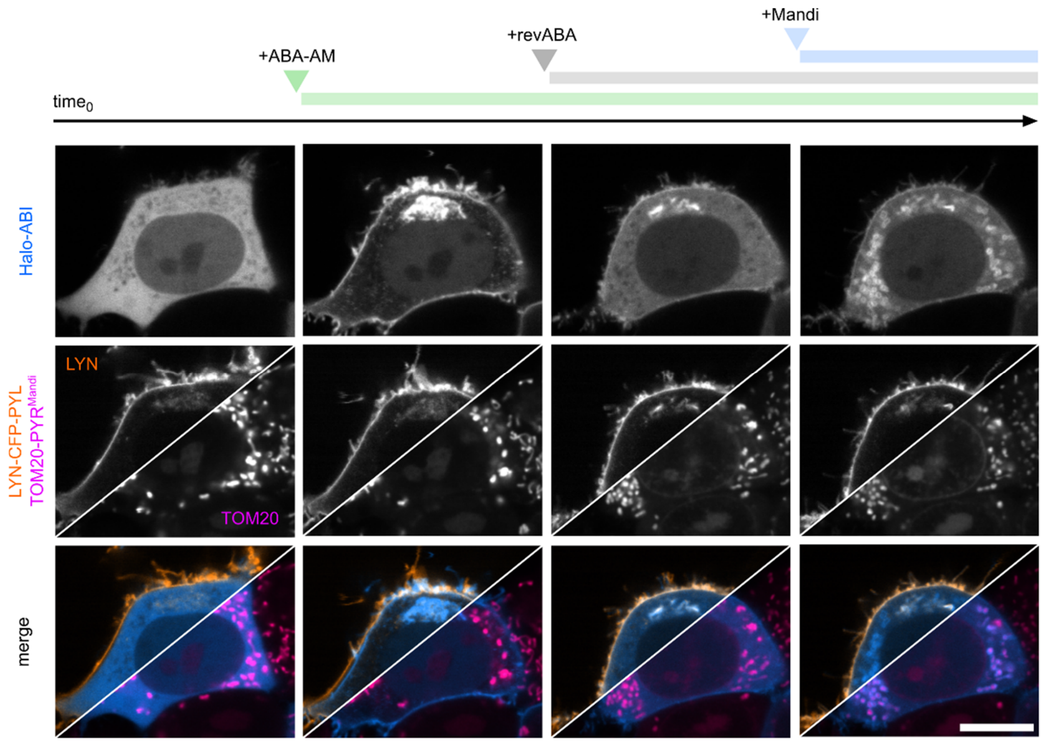


**Supplementary Figure 13: Reversible and dynamic protein shuttling between mitochondria and outer membrane in living cells.** HEK293T cells were co-transfected with TOM20-PYR$^{Mandi}$-IRES-Halo-ABI and LYN-CFP-PYL. Halo-tag domains was stained with HaloTag-SiR (20 nM) 2 h prior to imaging. Mitochondria were stained with MitoTracker Orange (100 nM, Thermo Fisher Scientific, USA) according to supplier protocol. Upper row shows dynamic receiver localization, middle row receptor localizations as references, lower row respective merges. Images acquired at $t_0$, 10 min after addition of ABA-AM (5 µM), 25 min after addition of revABA (100 µM), 10 min after addition of Mandi (5 µM). Scale bar 10 µm. Representative data for n>20 cells in 2 independent experiments.

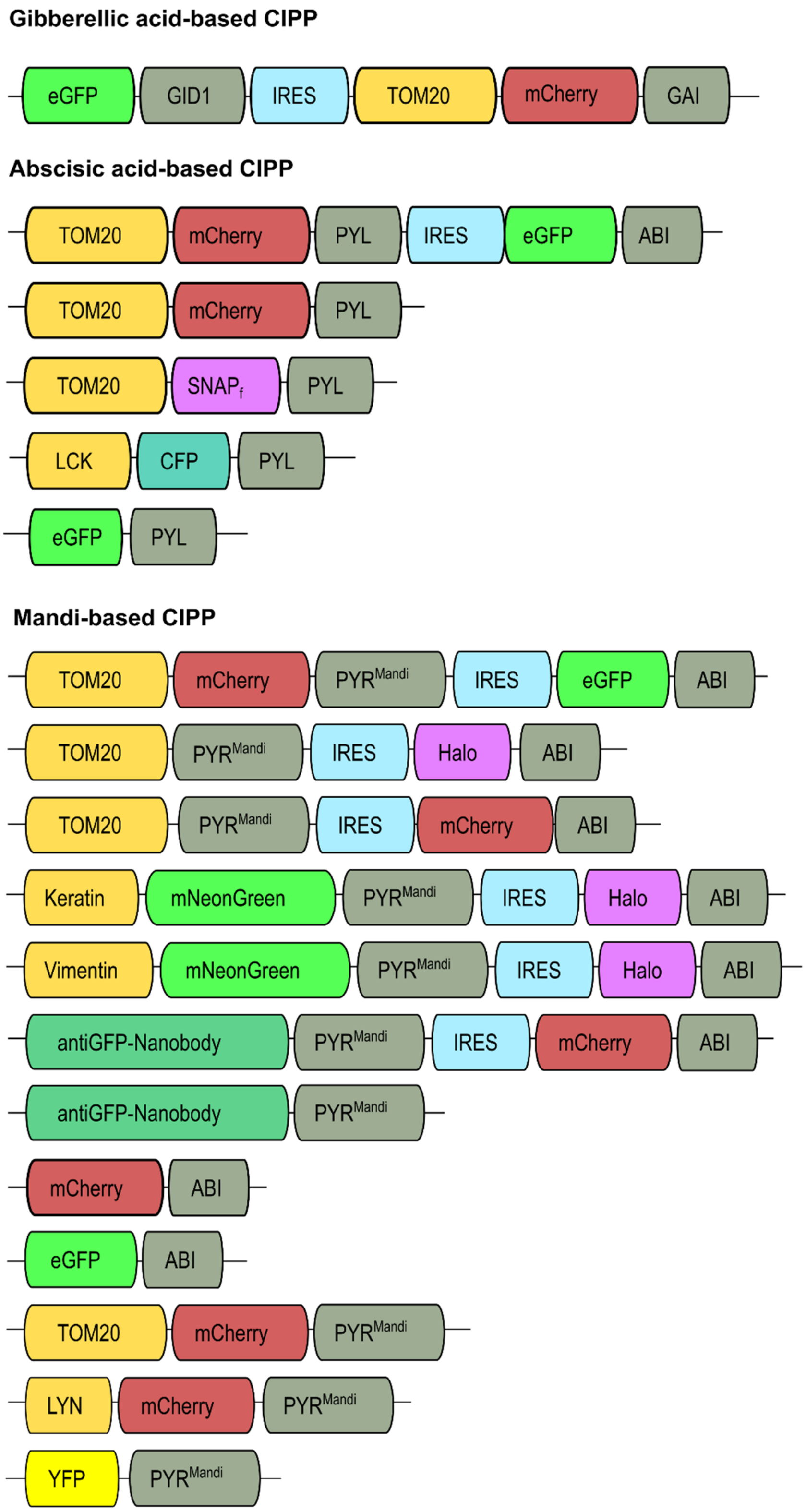


**Supplementary Figure 14: List of plasmids.**

### 9 Supplementary videos

**Supplementary video 1: Translocation of cytosolic eGFP-ABI to mitochondria-localized TOM20-mCherry-PYR$^{Mandi}$ after addition of Mandi to 100 nM final concentration.** COS-7 were transiently transfected with TOM20-mCherry-PYR$^{Mandi}$-IRES-EGFP-ABI. Epifluorescence timelapse images were acquired after addition of Mandi at t=0 sec. Raw data was background subtracted, flatfielded to correct for Gaussian-shaped illumination profile and temporally smoothed with a 2 frame running average. Scale bar: 10 µm. Representative data for 6 cells.

**Supplementary video 2: Translocation of cytosolic eGFP-ABI to mitochondria-localized TOM20-mCherry-PYR$^{Mandi}$ after addition of Mandi to 10 nM final concentration.** COS-7 were transiently transfected with TOM20-mCherry-PYR$^{Mandi}$-IRES-EGFP-ABI. Epifluorescence timelapse images were acquired after addition of Mandi at t=0 sec. Image pairs with 488 nm and 561 nm illumination for eGFP (green) and mCherry (magenta) excitation respectively were acquired for each time point. Raw data was background subtracted, flatfielded to correct for Gaussian-shaped illumination profile and temporally smoothed by a 2 frame running average projection. Scale bar: 10 µm. Representative data for 12 cells from n=2 independent experiments.

**Supplementary video 3: Translocation of cytosolic eGFP-ABI to plasma-membrane-localized LYN-mCherry-PYR$^{Mandi}$ in vivo.** eGFP-ABI and LYN-mCherry-PYR$^{Mandi}$ plasmids were injected in the york of 1-2 cell zebrafish embryos. Recruitment of eGFP to plasma membrane after addition of Mandi (final concentration 500 nM) at t=0 min, followed by confocal microscopy. Raw data was corrected for photobleaching and temporally smoothed by a 2 frame running average projection. Scale bar: 40 µm. Data representative for n=3 independent experiments.

**Supplementary video 4: Recruitment of ABI to PYL can be efficiently reversed using revABA.** COS-7 cells were transiently transfected with TOM20-mCherry-PYL-IRES-eGFP-ABI. EGFP-ABI was recruited to mitochondria by incubation with 200 nM ABA for 2 hours. Directly before imaging, medium was replaced with L15 containing 200 nM ABA. revABA at a concentration of 10 µM was then added during imaging at time point t=0 min and eGFP-ABI localization was followed over time. Scale bar: 10 µm. Representative data for 20 cells from n=3 independent experiments.

### 10 Materials

#### 10.1 Synthesis

##### 1.1.1 Mandipropamid

There are multiple sources of mandipropamid. For example, it can be purchased as pure compound from common suppliers or synthesized according to literature[17]. Here we present an alternative method to access gram scale: the extraction and isolation of mandipropamid from Revus TOP®, an agrochemical marketed by Syngenta.

###### *1.1.1.1 Extraction of Mandipropamid from Revus TOP®*

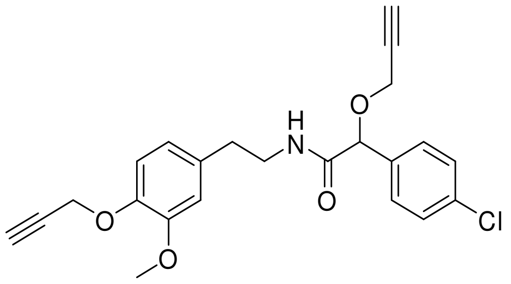


Mandipropamid was isolated from the commercially available suspension concentrate Revus TOP®, which contains mandipropamid and difenoconazol. 10 g Revus TOP® were suspended in 50 ml $H_2O$ and extracted 10-fold with 50 ml DCM. The yellow extract was dried over $Na_2SO_4$ and concentrated under reduced pressure. The crude product was purified by column chromatography ($SiO_2$, DCM:EA- 9:1) yielding in an off-white solid (2.2 g, 5.34 mmol).

$^1$H NMR (300 MHz, $CDCl_3$) δ 7.37 – 7.26 (m, 4H), 6.97 (d, *J* = 7.9 Hz, 1H), 6.70 (m, 3H), 4.96 (s, 1H), 4.78 – 4.74 (m, 2H), 4.11 (m, 2H), 3.83 (s, 3H), 3.54 (m, 2H), 2.79 (t, *J* = 6.7 Hz, 2H), 2.51 (t, *J* = 2.4 Hz, 1H), 2.48 (t, *J* = 2.4 Hz, 1H).

$^{13}$C NMR (75 MHz, $CDCl_3$) δ 169.5 (s, 1C), 149.8 (s, 1C), 145.4 (s, 1C), 134.7 (s, 1C), 134.6 (s, 1C), 132.7 (s, 1C), 128.8 (s, 2C), 128.7 (s, 2C), 120.6 (s, 1C), 114.7 (s, 1C), 112.4 (s, 1C), 79.7 (s, 1C), 75.8 (s, 1C), 75.7 (s, 1C), 56.9 (s, 2C), 56.4 (s, 2C), 55.7 (s, 1C), 40.2 (s, 1C), 35.2 (s, 1C).

HR-ESI$^{+:}$ m/z calcd. for $[C_{23}H_{22}ClNO_4+Na]^+$: 434.1130; found: 434.1123.

##### 1.1.2 Synthesis of acetoxymethyl substituted CIP

*1.1.2.1 (+)-Abscisic acid acetoxymethyl ester (ABA-AM)*

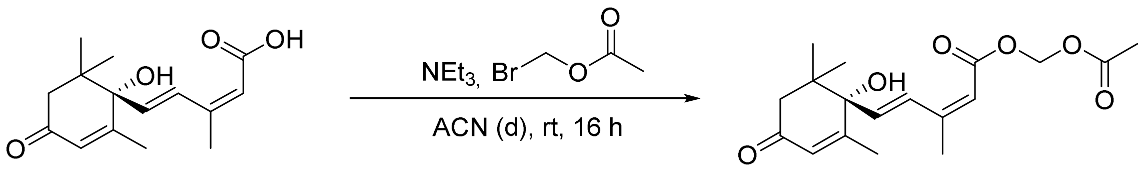


A flame-dried Schlenk tube was charged with (+)-abscisic acid (10.8 mg, 40.8 µmol, 1 eq) dissolved in 1 ml anhydrous acetonitrile. After addition of bromomethyl acetate (5.2 µl, 8.1 mg, 53.0 µmol, 1.3 eq) and triethylamine (7.0 µl, 5.4 mg, 53.0 µmol, 1.3 eq) the reaction mixture was stirred overnight at room temperature. The crude product was concentrated under reduced pressure and purified by column chromatography ($SiO_2$, Cy:EA- 2:1) yielding in an off-white resin (8.8 mg, 28.1 µmol, 69 %).

$^1$H NMR (500 MHz, $CD_3OD$) δ 7.79 (dd, J = 16.1, 0.9 Hz, 1H), 6.34 (dd, J = 16.1, 0.7 Hz, 1H), 5.94 (t, J = 1.3 Hz, 1H), 5.76 – 5.75 (m, 1H), 5.75 (s, 2H), 2.53 (d, J = 16.9 Hz, 1H), 2.20 (d, J = 16.4 Hz, 1H), 2.11 – 2.04 (m, 6H), 1.93 (s, 3H), 1.07 (s, 3H), 1.02 (s, 3H).

$^{13}$C NMR (126 MHz, $CDCl_3$) δ 197.7 (s, 1C), 170.1 (s, 1C), 164.3 (s, 1C), 162.3 (s, 1C), 152.1 (s, 1C), 137.6 (s, 1C), 128.0 (s, 1C), 127.3 (s, 1C), 117.2 (s, 1C), 79.8 (s, 1C), 79.1 (s, 1C), 49.9 (s, 1C), 41.7 (s, 1C), 24.5 (s, 1C), 23.2 (s, 1C), 21.5 (s, 1C), 21.0 (s, 1C), 19.0 (s, 1C)

HR-ESI$^{+}$: m/z calcd. for $[C_{18}H_{24}O_6+Na]^+$: 359.1465; found: 359.1477.

*1.1.2.2 Gibberellic acid acetoxymethyl ester ($GA_3$-AM)*

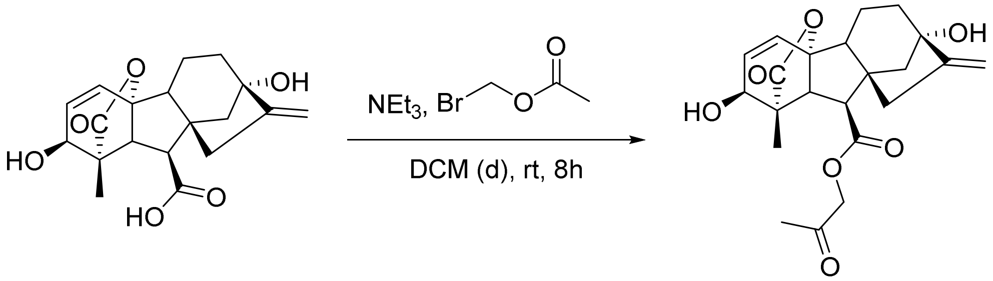


$GA_3$-AM was prepared according to procedures described in previous publications[18].

##### 1.1.3 Synthesis of abscisic acid antagonist revABA

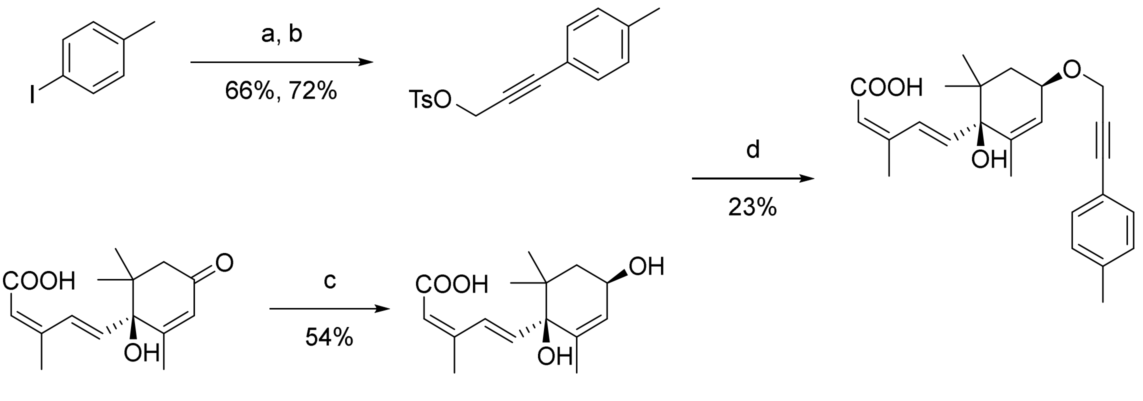


**Scheme S 1 Route for synthesis of ABA antagonist revABA[19]. a,** propargylalcohol, $(Ph_3P)_2PdCl_2$, CuI, DIPEA, THF; **b,** TsCl, KOH, Et2O; **c:** NaBH4, $CeCl_3$*$7H_2O$, MeOH **d,** NaH, THF.

###### *1.1.3.1 (p-Tolyl)prop-2-yn-1-ol*

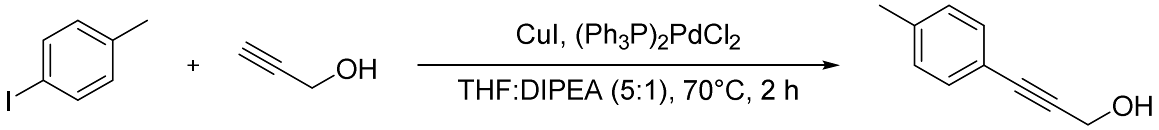


*p*-Iodo toluene (1.0 g, 4.6 mmol, 1 eq) was dissolved in a mixture of 5 ml THF and 1 ml DIPEA. After addition of propargyl alcohol (300 µl, 5.50 mmol, 1.2 eq) the mixture was degassed by bubbling argon through the solution for 10 min. After addition of copper(I) iodide (43.7 mg, 229.3 µmol, 0.05 eq) and bis(triphenylphosphine)palladium(II) dichloride (64.4 mg, 91.7 µmol, 0.02 eq) the reaction mixture was heated to 70°C and stirred for 2 h. The reaction was quenched with a 1 M aqueous solution of HCl and extracted three times with DCM. The organic layer was dried over $Na_2SO_4$ and concentrated under reduced pressure. The crude product was purified by column chromatography ($SiO_2$, Cy:EA- 9:1) yielding a brown oil (446.8 mg, 3.06 mmol, 67%). The measured NMR spectrum was in accordance with literature reported spectra[19].

$^1H$ NMR (300 MHz, $CDCl_3$) δ 7.33 (d, *J* = 8.0 Hz, 2H), 7.12 (d, *J* = 8.0 Hz, 2H), 4.49 (d, *J* = 6.1 Hz, 2H), 2.34 (s, 3H), 1.65 (t, *J* = 6.1 Hz, 1H).

$^{13}C$ NMR (75 MHz, $CDCl_3$) δ 138.23 (s, 1C), 131.15 (s, 2C), 128.64 (s, 2C), 118.98 (s, 1C), 86.04 (s, 1C), 85.45 (s, 1C), 51.31 (s, 1C), 21.05 (s, 1C).

*1.1.3.2 3-(p-Tolyl)prop-2-yn-1-yl 4-methylbenzenesulfonate*

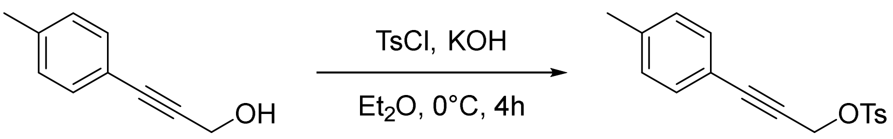


According to Omar *et al.*[20], tosyl chloride (26.1 mg, 136.8 µmol, 1 eq) and (*p*-tolyl)prop-2-in-1-ol (20.0 mg, 136.8 µmol, 1 eq) were dissolved in 5 ml diethyl ether and cooled to 0 °C. Freshly grinded KOH (76.8 mg, 1,37 mmol, 10 eq) was slowly added and the reaction was stirred for further 4 h at 0 °C. After TLC showed full conversion, the reaction mixture was quenched with ice water and extracted with diethyl ether. The combined organic layer was dried over $Na_2SO_4$ and concentrated under reduced pressure. The product was obtained as a brown oil (29.9 mg, 99.5 µmol, 73%). The measured NMR spectrum was in accordance with literature and the product was directly used without further purification.

$^1$H NMR (300 MHz, Chloroform-*d*) δ 7.85 (d, *J* = 8.1 Hz, 2H), 7.31 (d, *J* = 8.0 Hz, 2H), 7.15 (d, *J* = 8.1 Hz, 2H), 7.08 (m, *J* = 8.0 Hz, 2H), 4.94 (s, 2H), 2.40 (s, 3H), 2.34 (d, *J* = 3.4 Hz, 3H).

$^{13}$C NMR (126 MHz, Chloroform-d) δ 144.9 (s, 1C), 139.3 (s, 1C),133.4 (s, 1C), 131.7 (s, 2C), 129.8 (s, 2C), 129.0 (s, 2C), 128.2 (s, 2C), 118.3 (s, 1C), 89.2 (s, 1C), 79.9 (s, 1C), 58.8 (s, 1C), 21.6 (s, 1C), 21.5 (s, 1C).

HR-ESI$^{+:}$ m/z calcd. for $[C_{17}H_{16}O_3S+Na]^+$: 323.0718; found: 323.0724.

*1.1.3.3 ABA-(S)-OH*

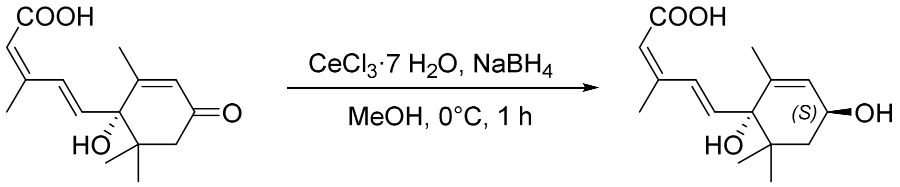


According to Takeuchi *et al.*[19], a flame-dried Schlenk flask was equipped with abscisic acid (50 mg, 189.2 µmol, 1 eq) and dissolved in 4 ml dry methanol. The solution was cooled to 0°C and cerium(III) chloride heptahydrate (211.4 mg, 567.5 µmol, 3 eq) and sodium borohydride (15.1 mg, 399.1 µmol, 2.1 eq) were added. The reaction mixture was stirred for further 60 min and quenched with a saturated solution of ammonium chloride. After concentration of the solution in vacuo to remove methanol, the reaction mixture was acidified to pH 2 by addition of aqueous 1 M HCl and extracted with DCM. The combined organic layer was washed with brine, dried over $Na_2SO_4$ and concentrated under reduced pressure. The crude product was purified by column chromatography ($SiO_2$, Cy:EA- 3:1), yielding in a colorless, cloudy oil (31.3 mg, 117.5 µmol, 62 %).

$^1$H NMR (500 MHz, Methanol-*d4*) 7.79 – 7.60 (m, 1H), 6.21 (d, J = 16.1 Hz, 1H), 5.69 (d, J = 1.6 Hz, 1H), 5.52 (dt, J = 2.8, 1.4 Hz, 1H), 4.18 (ddt, J = 10.4, 6.4, 2.2 Hz, 1H), 2.01 (d, J = 1.3 Hz, 3H), 1.71 (m, 1H), 1.64 (s, 3H), 1.63 – 1.55 (m, 1H), 1.01 (s, 3H), 0.90 (s, 3H).

$^{13}C$ NMR (126 MHz, Methanol-*d4*) δ 169.7 (s, 1C), 152.0 (s, 1C), 141.4 (s, 1C), 139.5 (s, 1C), 128.8 (s, 1C), 128.0 (s, 1C), 118.4 (s, 1C), 80.3 (s, 1C), 66.4 (s, 1C), 44.9 (s, 1C), 41.0 (s, 1C), 25.6 (s, 1C), 23.2 (s, 1C), 21.4 (s, 1C), 18.2 (s, 1C).

HR-ESI$^{+:}$ m/z calcd. for $[C_{15}H_{22}O_4+Na]^+$: 289.1416; found: 289.1423.

*1.1.3.4 ABA antagonist revABA*

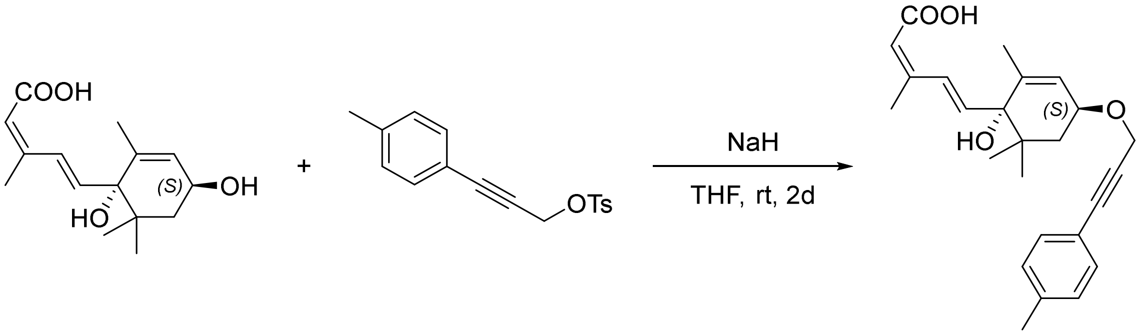


According to literature[19], a flame-dried Schlenk tube was equipped with sodium hydride (10.2 mg, 60 wt%, 255.3 µmol, 3.4 eq), which was suspended in 5 ml dry THF, cooled to 0°C. ABA-(*S*)-OH (20.0 mg, 75.1 µmol, 1 eq) was added and the solution was stirred for 15 min. In a separate flame-dried Schlenk tube, 3-(*p*-Tolyl)prop-2-in-1-yl 4-methylbenzolsulfonat (28.2 mg, 80 wt%, 75.1 µmol, 1 eq))was dissolved in 2 ml dry THF and slowly added to the reaction mixture. The reaction was allowed to warm up to room temperature and stirred for 2 d under an argon atmosphere. As the reaction did not show full conversion according TLC, another 6.8 eq NaH (20.4 mg, 0.51 mmol) were added to the reaction mixture and the reaction was stirred for further 16 h at 50°C. The reaction was quenched with 1 M HCl solution and extracted with DCM. The organic layer was washed with brine, dried over $Na_2SO_4$ and concentrated under reduced pressure. The product was purified by column chromatography ($SiO_2$, Cy:EA 0-50%) and obtained as a yellowish resin (6.8 mg, 17.2 µmol, 23%). For live cell experiments a batch of revABA was further purified by preparative reverse phase HPLC (C18, $H_2O$/MeCN gradient +0.1% TFA).

$^{1}H$ NMR (500 MHz, DMSO-*d6*) δ 7.62 (d, *J* = 15.9 Hz, 1H), 7.32 (d, *J* = 7.8 Hz, 2H), 7.19 (d, *J* = 7.8 Hz, 2H), 6.08 (d, *J* = 15.9 Hz, 1H), 5.63 (s, 1H), 5.57 (s, 1H), 4.39 (d, *J* = 1.5 Hz, 2H), 4.13 (m, 1H), 2.31 (s, 3H), 1.94 (d, *J* = 1.1 Hz, 3H), 1.78 (m, 1H) 1.57 (m, 4H), 0.93 (s, 3H), 0.84 (s, 3H).

$^{13}C$ NMR (126 MHz, DMSO-*d6*) δ 166.9 (s, 1C), 149.4 (s, 1C), 140.3 (s, 1C), 139.2 (s, 1C), 138.4 (s, 1C), 131.2 (s, 1C), 129.3 (s, 2C), 125.9 (s, 2C), 123.8 (s, 1C), 119.0 (s, 1C), 117.7 (s, 1C), 86.1 (s, 1C), 85.2 (s, 1C), 77.6 (s, 1C), 71.9 (s, 1C), 55.3 (s, 1C), 40.1 (s, 1C), 39.3 (s, 1C), 25.2 (s, 1C), 22.9 (s, 1C), 20.9 (s, 2C), 17.9 (s, 1C).

HR-ESI$^{+:}$ m/z calcd. for $[C_{25}H_{30}O_4+Na]^+$: 417.2042; found: 417.2041.

### 11 NMR Characterization

#### 11.1 Mandipropamid

$^{1}H$ NMR

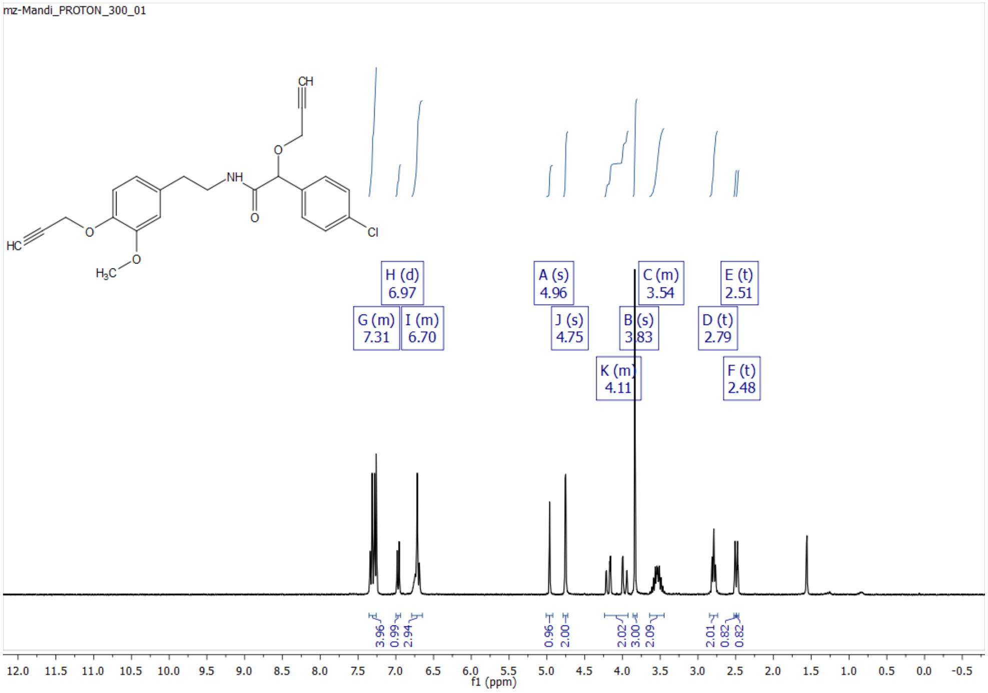


$^{13}C$ NMR/APT

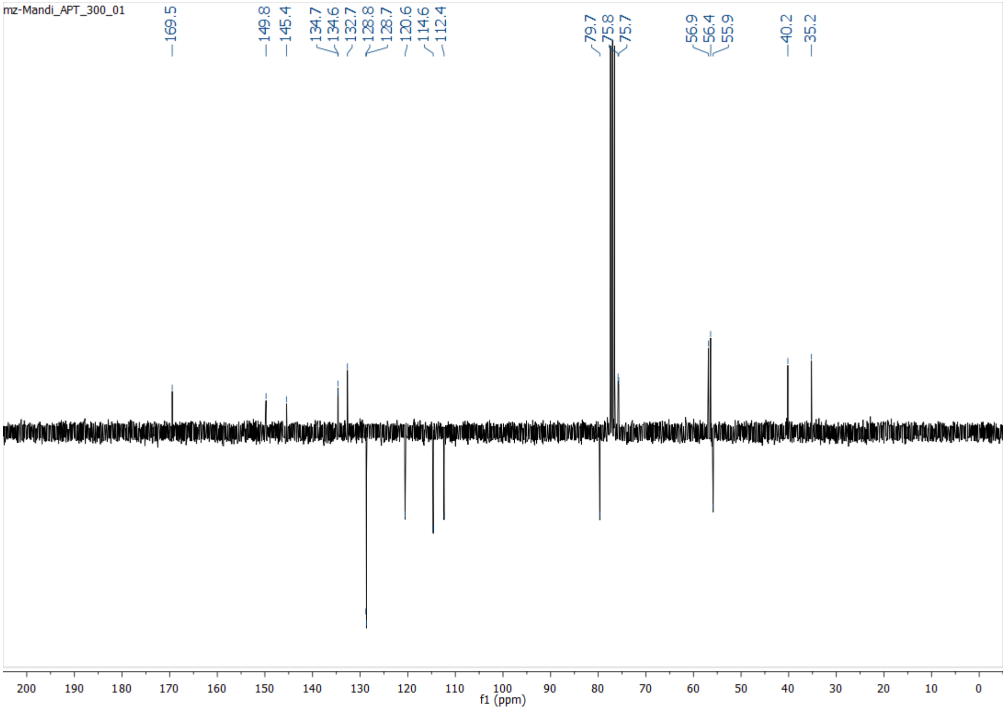

#### 11.2 ABA-AM

$^{1}$H NMR

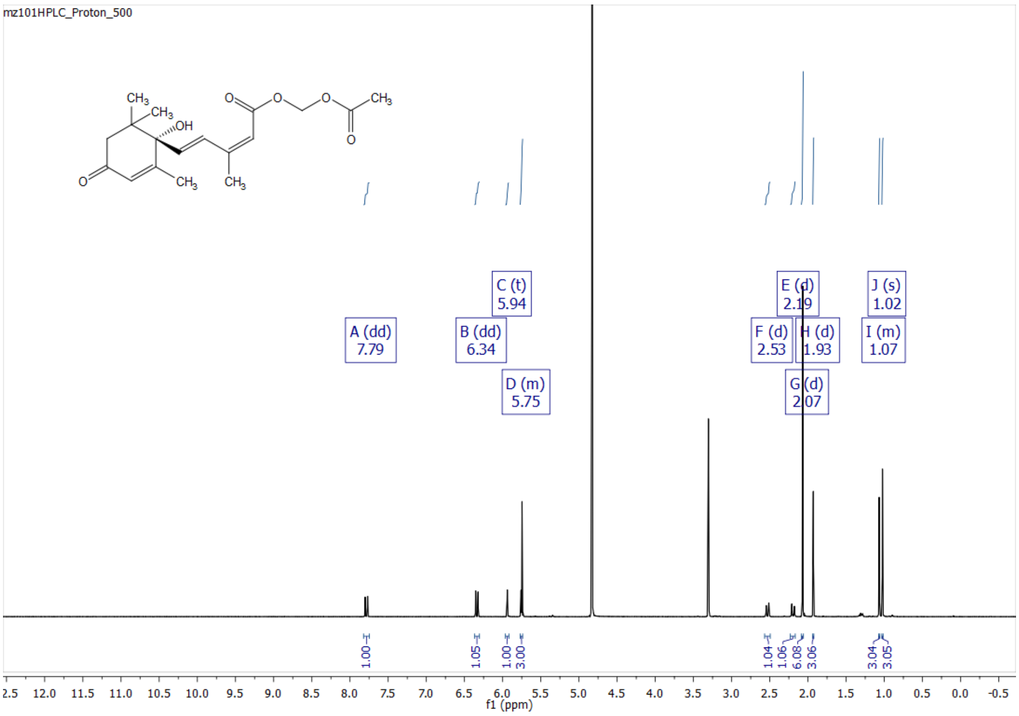


$^{13}$C NMR/APT

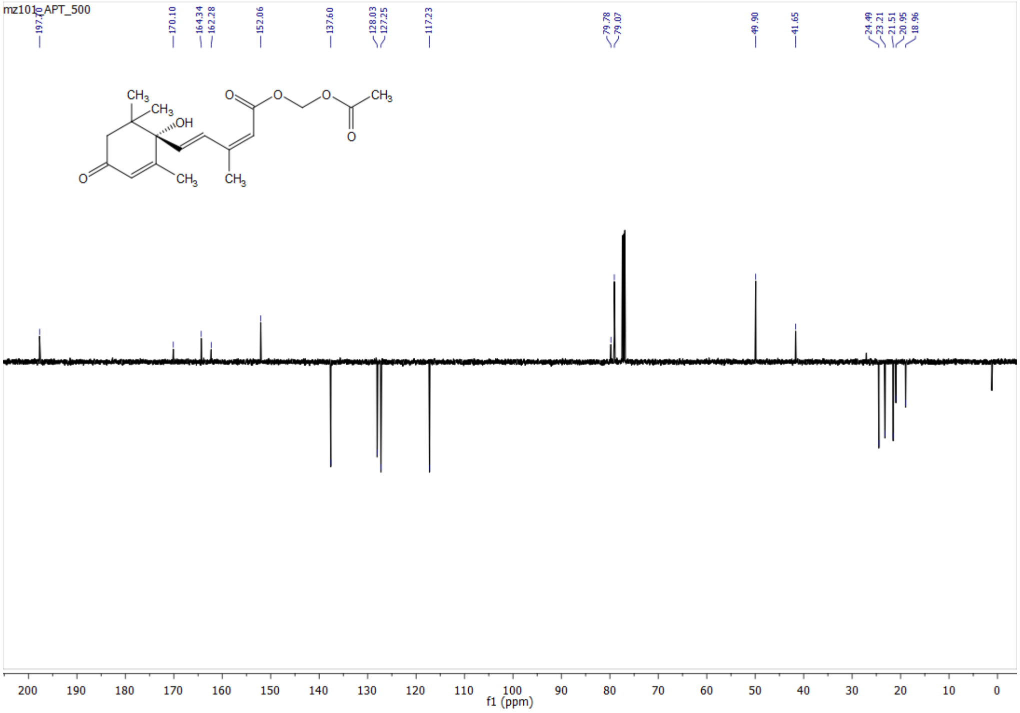

#### 11.3 (p-Tolyl)prop-2-yn-1-ol

$^{1}$H NMR

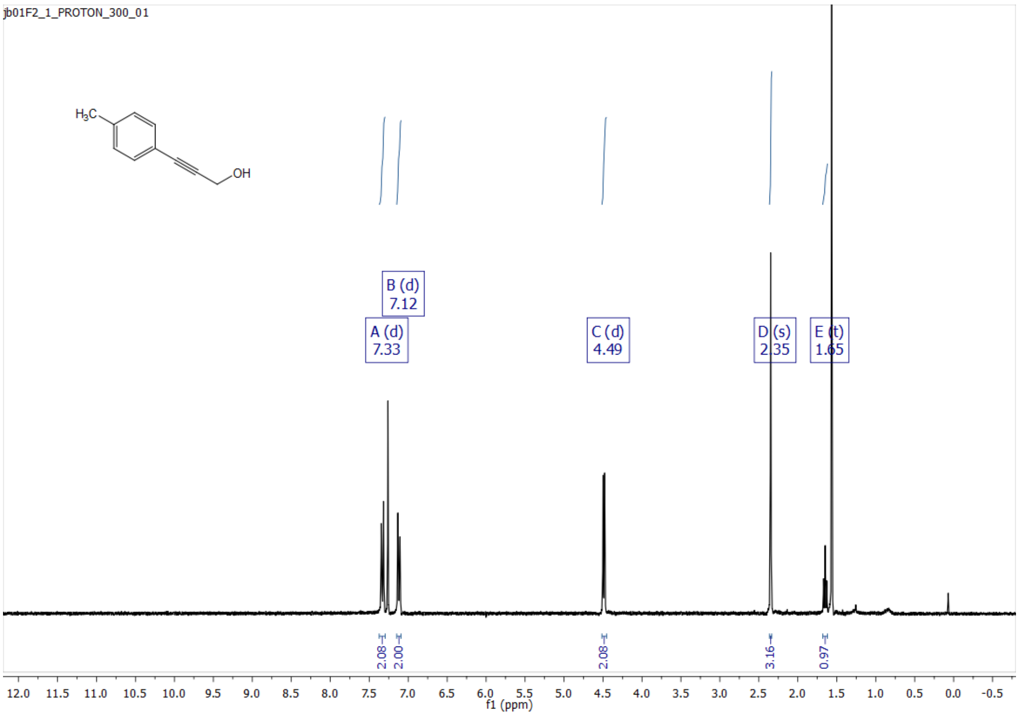


$^{13}$C NMR/APT

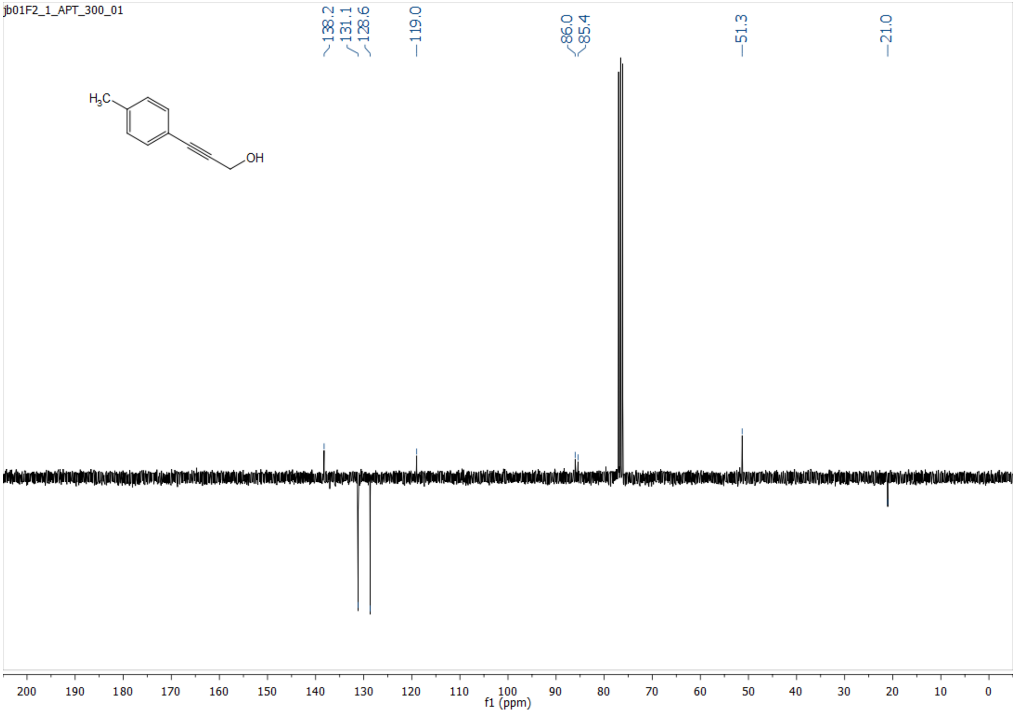

#### 11.4 3-(p-Tolyl)prop-2-yn-1-yl 4-methylbenzolsulfonate

$^{1}$H NMR

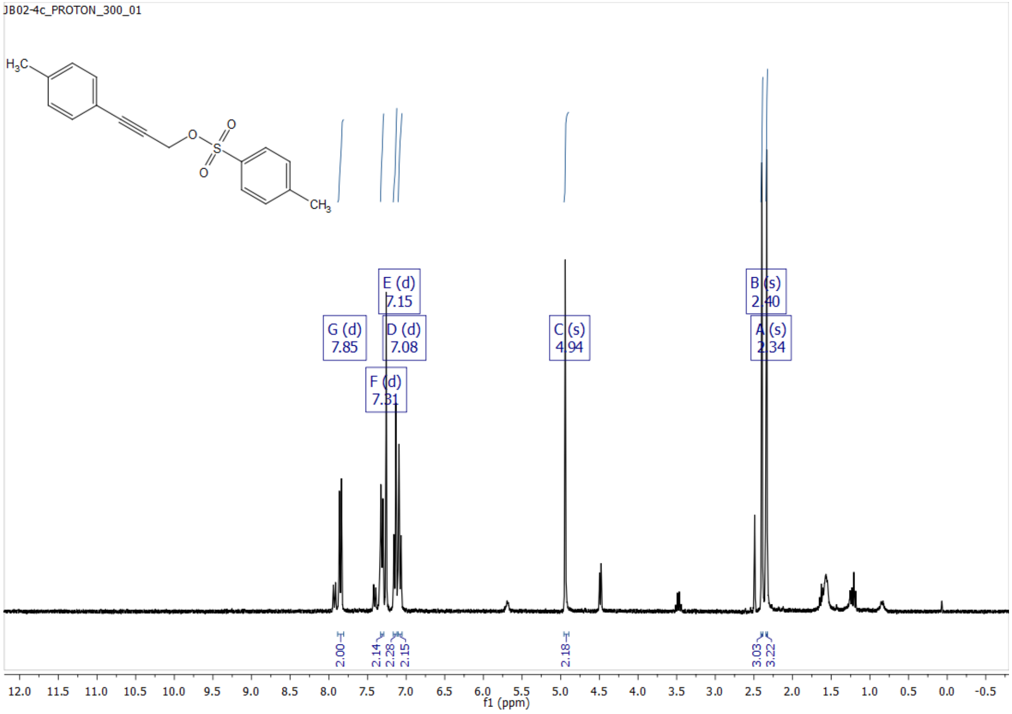


$^{13}$C NMR/APT

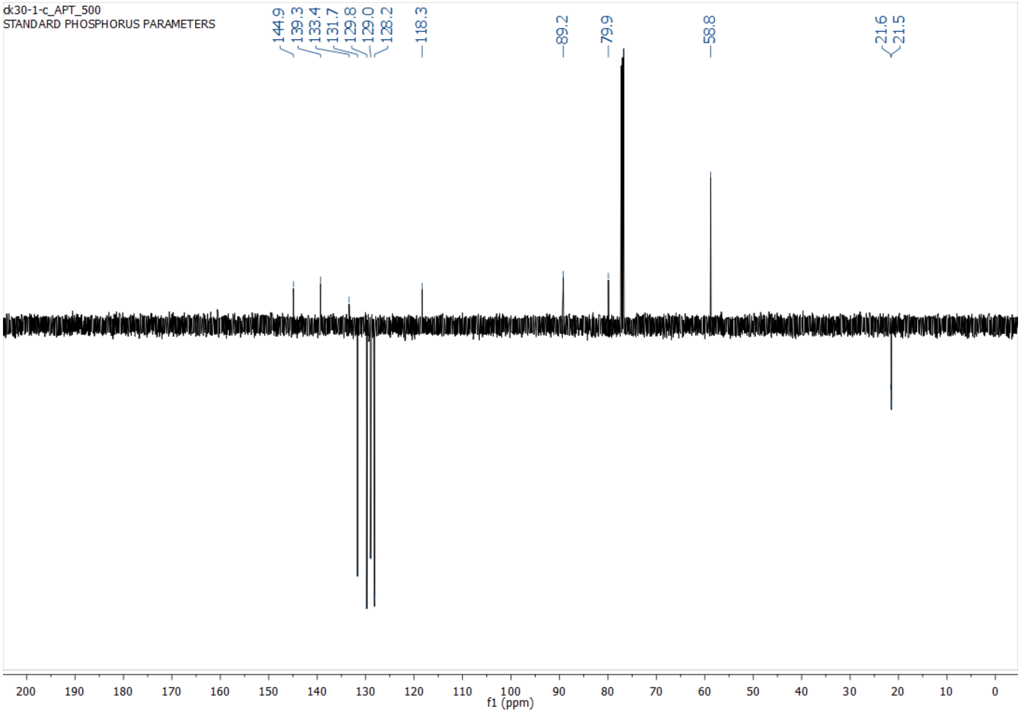

#### 11.5 ABA-(*S*)-OH

[1]H NMR

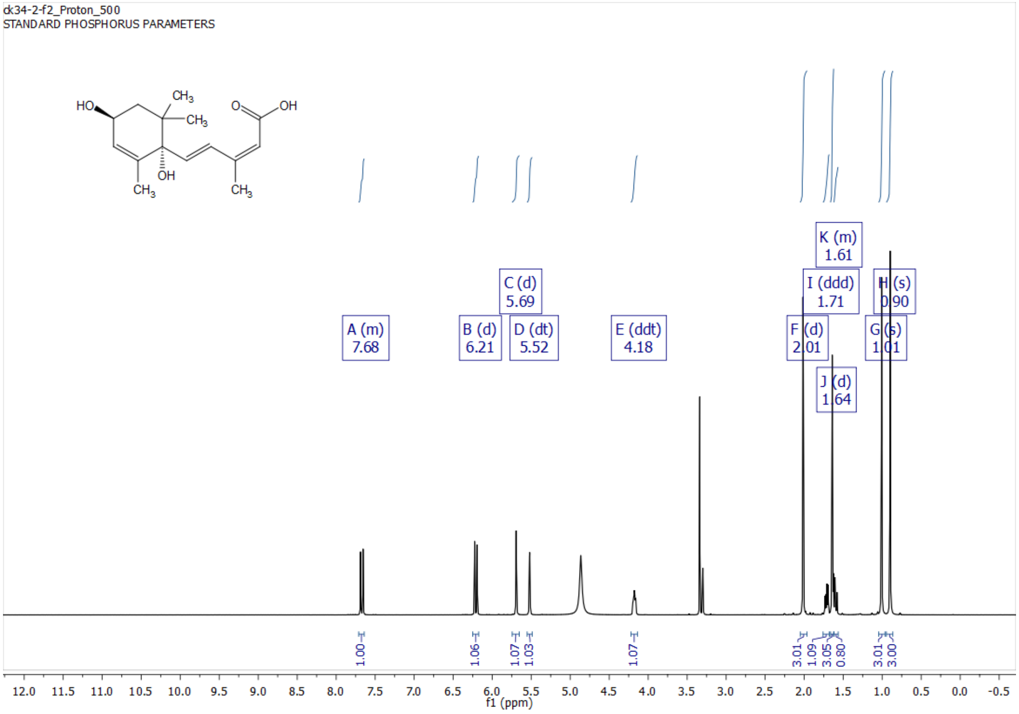


[13]C NMR-APT

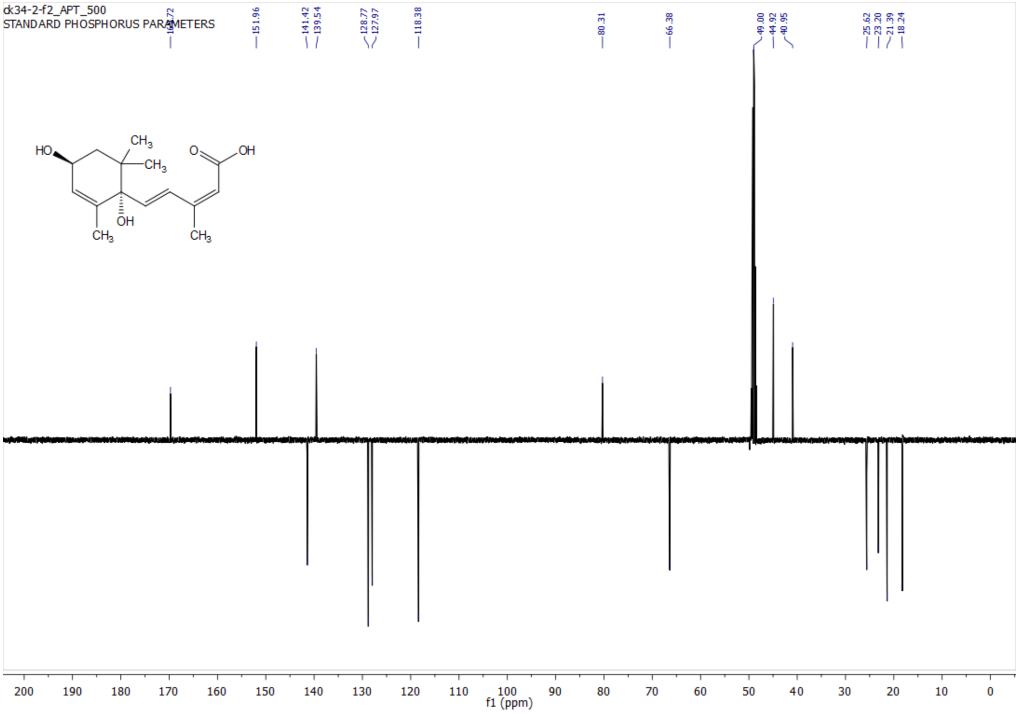

#### 11.6 revABA

$^{1}$H NMR

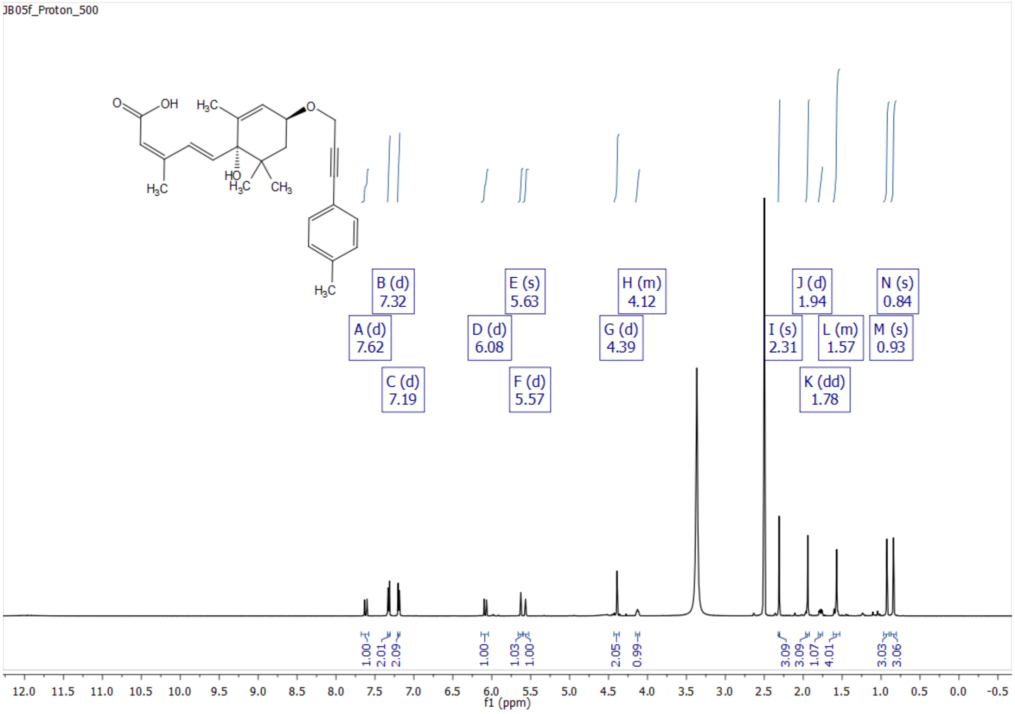


$^{13}$C NMR-APT

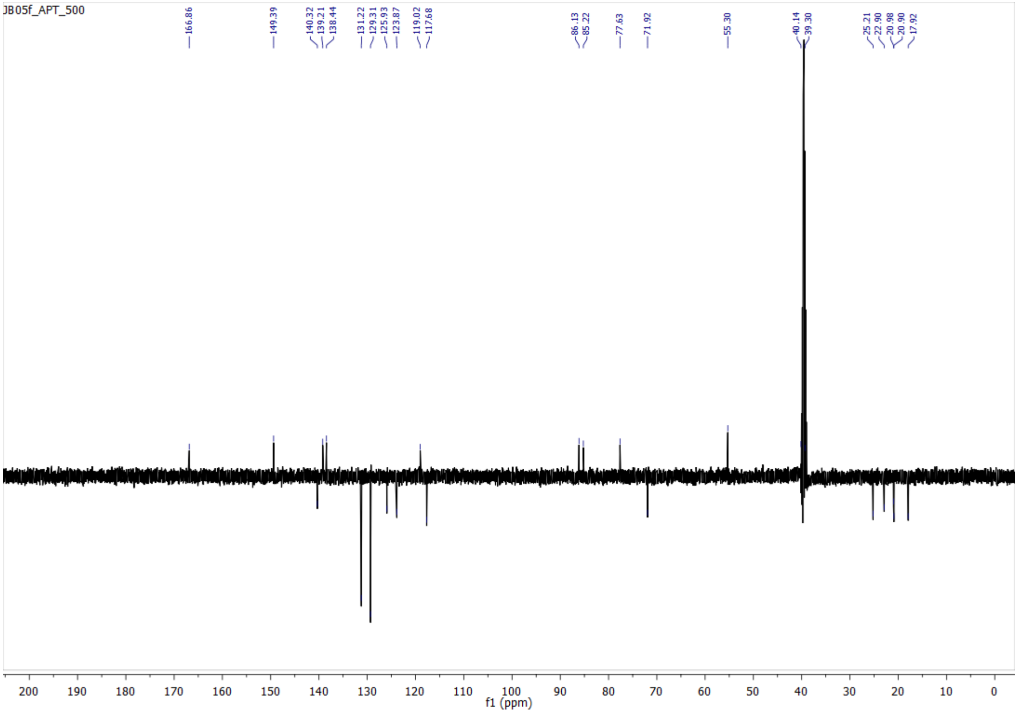
